# High-Sensitivity Top-Down Proteomics Reveals Enhanced Maturation of Micropatterned Induced Pluripotent Stem Cell-Derived Cardiomyocytes

**DOI:** 10.1101/2025.05.12.653499

**Authors:** Mallory C. Wilson, Mitchell Josvai, Janay K. Walters, Jodi Lawson, Kalina J. Rossler, Zhan Gao, Yanlong Zhu, Timothy J. Kamp, Wendy C. Crone, Lee L. Eckhardt, Ying Ge

## Abstract

Human induced pluripotent stem cell-derived cardiomyocytes (hiPSC-CMs) are increasingly used for disease modeling, drug discovery, and precision medicine, yet their utility is often limited by their immature phenotype. One promising maturation strategy involves using micropatterned substrates that mimic native cardiomyocytes’ organizational growth and stiffness. However, the molecular maturity of this model has not yet been assessed, and there is currently no method to extract proteins from micropatterned hiPSC-CMs for top-down proteomic analysis. Herein, we present a high-sensitivity, surfactant-free protein extraction protocol for the top-down proteomic analysis of hiPSC-CMs. Through this method, we assessed the maturation of micropatterned hiPSC-CMs compared to traditional unstructured monoculture and co-culture monolayers at the proteoform level. We found that micropatterned hiPSC-CMs display molecular signatures of cardiomyocyte maturations including increased expression of ventricular myosin light chain isoforms, reduced expression of the fetal troponin T isoform, and decreased phosphorylation of alpha-tropomyosin. This surfactant-free, high-sensitivity approach enables robust top-down proteomics from limited, heterogeneous cell populations, and identifies the micropattern hiPSC-CM as a more adult-like CM model, broadening the utility of structured culture systems for cardiac disease modeling and translational research.

## INTRODUCTION

Over the last two decades, human induced pluripotent stem cell-derived cardiomyocytes (hiPSC-CMs) have rapidly grown in popularity for disease modeling, drug discovery, and precision medicine.^1-5^ By enabling a personalized “disease-in-a-dish” approach, these *in vitro* models have revolutionized cardiovascular research towards understanding cardiac diseases and assessing cardiotoxicity directly from patients’ own cells.^6-9^ Despite their high promise, hiPSC-CM models are still severely limited by their relative immaturity compared to adult cardiomyocytes. This is largely due to the difficulty in replicating complex *in vivo* maturation cues and the inherently slow cardiomyocyte maturation in humans.^10,11^ This results in fetal-like models that can lack the contractile properties, electrophysiology, metabolism, and protein expression necessary to accurately represent an adult patient’s disease phenotype.^3,5,9^

To promote better maturation in hiPSC-CM models, a variety of strategies have been explored to promote maturation, including metabolic, electrical, mechanical conditioning, and microengineered platforms.^12-14^ A promising strategy towards maturation is to culture hiPSC-CMs on micropatterned substrates that mimic biological conditions by aligning seeded cardiomyocytes along interconnected lanes to more accurately reflect their *in vivo* geometry than a traditional 2D monolayer.^15,16^ Recently, polydimethylsiloxane (PDMS)-based micropatterns have combined reduced substrate stiffness, chevron lanes, and co-culture alongside hiPSC-derived fibroblasts (hiPSC-CFs) to promote native-like development after just 30 days of culture time.^17,18^ However, the molecular maturity of hiPSC-CMs cultured on these micropatterned substrates remains poorly understood, leaving a gap in understanding the applicability of these models.

Proteoforms - the distinct molecular forms arising from genetic variation, alternative splicing isoforms, and post-translational modifications (PTMs) - play critical roles in defining cellular function and phenotype.^19-21^ Top-down proteomics,^21,22^ which analyzes intact proteoforms, provides an unbiased method to assess cardiomyocyte maturation by capturing changes in protein isoform expression and PTMs.^10,23,24^ Previous studies have identified these markers of maturation by examining sarcomeric protein transitions, particularly the switch from fetal to adult isoforms,^10,25,26^ but detecting these transitions with top-down proteomics has traditionally required a large number of hiPSC-CMs (>1 million cells).^10,23^ Micropatterned hiPSC-CM cultures yield limited cell numbers much lower than previous models (∼50,000 hiPSC-CMs),^16,17^ necessitating the development of sensitive top-down methods to enable the proteoform-level maturation assessment of these models.

Herein, we present a highly sensitive, surfactant-free top-down proteomics platform that enables robust assessment of hiPSC-CM maturation from limited cell numbers cultured on PDMS micropatterns. With this optimized protocol, we achieve top-down quantitative proteomics of sarcomeric proteoforms directly from micropatterned hiPSC-CMs, for the first time. Our findings reveal that micropatterning of a soft substrate markedly accelerates cardiomyocyte proteomic maturation, with isoform switching and changes in the relative abundance of proteoforms comparable to 3D cultures maintained for twice as long. This advancement establishes a robust strategy for hiPSC-CM proteomic analysis from limited cell input, enabling the assessment of functionally mature models with improved reproducibility and translational relevance for disease modeling and drug discovery.

## EXPERIMENTAL PROCEDURES

### hiPSC-CM Culture & Harvest

A full description of the hiPSC to hiPSC-CM differentiation protocol using the small molecule GiWi method^27^ can be found in the Supplemental Methods. After differentiation and Magnetic-Activated cell sorting (MACS, Miltentyi) purification (Day 16), hiPSC-CMs were split between monoculture monolayers (MM), co-culture monolayers (CC), and co-culture micropatterns (µP). MM hiPSC-CMs were cultured on PDMS-coated tissue culture plastic (TCP) and cultured until Day 30. CC hiPSC-CMs were seeded at a 10:1 ratio of cardiomyocytes to fibroblasts on PDMS-coated TCP and cultured for14 days (Day 30 total culture time). µP hiPSC-CMs were seeded at a 10:1 ratio of cardiomyocytes to fibroblasts on micropatterned PDMS and cultured until Day 14 (Day 30 total culture time). Cells were harvested and singularized by TrypLE 10X (Life Technologies) incubation for 15 minutes followed by an immediate trypsin quench via dilution with EB20 medium (*Supplemental Methods*). To isolate hiPSC-CMs, heterogenous solubilized cells were incubated on uncoated TCP for 60 min at 37°C. The supernatant was removed, enabling a pseudo-enrichment of hiPSC-CMs and leaving hiPSC-CFs attached to the uncoated TCP. Though hiPSC-CFs were not present in the MM cohort, those samples were also subjected to a 60 min incubation and subsequent supernatant removal to ensure treatment consistency. Isolated hiPSC-CM samples were centrifuged at (1000 rpm, 10 min, 25°C), then washed with Dulbecco’s phosphate-buffered saline (DPBS, Thermo Fisher Scientific). DPBS-solubilized monolayer samples were aliquoted to 50k cells/sample to match individual micropattern cell counts. All samples were moved to Protein LoBind centrifuge tubes (Eppendorf) and cells were pelleted via centrifugation (21,000 × g, 15 min, 4 °C). The supernatant was removed, and the samples were stored at -80°C until protein extraction.

### Surfactant-free Protein Extraction

The protein extraction protocol was inspired by the hexafluoroisopropanol (HFIP)-enabled single muscle fiber extraction.^28^ On the day of mass spectral analysis, cell pellets were thawed (∼1 min) and resuspended in 90% HFIP extraction solution (90% HFIP, 10 mM L-methionine, 1× HALT protease & phosphatase inhibitor cocktail) via gentle up-and-down pipetting before a 15 min incubation on ice. Samples were then diluted with 20 µL of 0.2% formic acid (FA) and briefly dipped (<2 sec) into an ultrasonic water bath three times, followed by three freeze-thaw cycles (incubation at −80 °C for 10 min followed immediately by incubation for 1 min at 37 °C). Gentle agitation was applied between each freeze-thaw cycle. Samples were centrifuged (21,000 × *g*, 15 min, 4 °C), and the supernatant was removed for desalting via buffer exchange. Buffer exchange with 0.2% FA was performed using a 10 kDa molecular weight cutoff spin filter, then samples were further concentrated to a final volume of around 35 µL. Sample concentrations were quantified using a standard Bradford assay, and the remaining lysate was transferred to an HPLC vial.

### Liquid Chromatography & Mass Spectrometry

OtofControl 3.4 (Bruker Daltonics) was used to collect all LC-MS data. Protein separation was performed using reverse-phase chromatography with a NanoAcuity Ultra-High Pressure LC system (Waters, Milford, MA, USA). Injection volumes were adjusted to enable 500 ng of total protein. Elution was performed from a home-packed C4 column (200 × 0.250 mm, 2.7 µm, 1000 Å C4 (Halo)) held at 50°C with a flow rate of 3 µL/min. Mobile phase A (MPA) contained 0.2% FA in H_2_O, and mobile phase B (MPB) contained 0.2% FA in ACN, with a 65 min RPC gradient of the following MPB concentrations: start at 10% MPB, hold 10% until 5 min, 25% at 15 min, 40% at 35 min, 50% at 45 min, 95% at 55 min, adjusted back to 10% at 55.1 min, and held at 10% until 65 min. Eluted proteins were electrosprayed into a high-resolution Impact II quadrupole time-of-flight (QTOF) mass spectrometer (Bruker Daltonics, Bremen, Germany). The end plate offset was set to 500 V and the capillary voltage was set to 4500 V. The nebulizer was set to 0.5 bar with a dry gas flow rate of 4.0 L/min at 200°C. Mass spectra were collected at a scan rate of 1 Hz over 300-3000 m/z with a quadrupole low mass set to 650 m/z.

## Data Analysis

LC-MS data was processed and analyzed using DataAnalysis (v4.3, Bruker Daltonics) software.^29^ Maximum Entropy Deconvolution of spectra was performed with a resolving power of 60,000 for isotopically resolved proteins. The top 5-7 most abundant charge state ions from the non-deconvoluted spectra were used to produce extracted ion chromatograms (EICs), and relative abundances of isoforms were measured using the ratios of the area under the curve (AUC) of each isoform’s EIC. Relative quantification of proteoforms was performed by deconvoluting mass spectra using the Maximum Entropy Deconvolution algorithm and taking the ratio of the highest peak intensity of the proteoform to the summed intensities of all proteoforms of that protein. Total phosphorylation of proteins with multiple phosphorylation sites was calculated as the ratio of the sum of the peak intensity of all phosphorylated proteoforms (multiplied by the integer number of phosphorylated sites on that proteoform) to the sum of all proteoform peak intensities of that protein. The sophisticated numerical annotation procedure (SNAP) algorithm was applied to determine the monoisotopic masses of all observed ions. Normality was determined using the linearity of Q-Q plots and the Shapiro-Wilke test. Statistical analysis was performed using a one-way Analysis of Variance (ANOVA) to compare the means of all three cohorts. Post hoc analysis was performed using Tukey’s Honest Significance Difference (HSD) test for pairwise comparisons; “n.s.” indicates a *p* > 0.05, * indicates *p* < 0.05, and ** indicates a *p* < 0.001.

## RESULTS AND DISCUSSION

### High-sensitivity top-down proteomics for micropatterned hiPSC-CMs enabled by surfactant-free extraction

First, we aim to establish a high-sensitivity, surfactant-free protein extraction method for a samples with limited numbers of hiPSC-CMs inspired by our previously developed high-sensitivity top-down proteomics method for single muscle cells.^28^ The original surfactant-free protein extraction protocol successfully extracted proteins from skeletal muscle cells using just 25% hexafluoroisopropanol (HFIP), a weakly acidic fluoroalcohol that clusters similarly to surfactants in aqueous solutions.^30^ However, it did not provide efficient extraction of proteins from hiPSC-CM samples, potentially due to differences in the membrane composition, extracellular matrix laid during time in culture, and intercellular junctions, like intercalated discs.^31^ To optimize the extraction for the micropattern platform, we increased the concentration of HFIP to 90% and incorporated pulse sonication by brief sample tube immersion in a sonicating water bath (*Supplementary Figure S1*). Though these strategies alone were not effective solutions, their combination achieved robust cell lysis and solubilization of proteins, facilitating the highly reproducible proteoform-level quantitative analysis of sarcomeric proteins from micropatterned hiPSC-CMs. This is demonstrated by consistent retention times and spectral intensities of three LC-MS technical replicates and consistent linear response of varied protein loading injection replicates (*Supplementary Figure S2*).

Next, we applied the optimized surfactant-free extraction protocol to hiPSC-CM samples cultured under three distinct conditions (n=8 per group): monoculture monolayers (MMs), co-culture monolayers with hiPSC-CFs (CCs), and co-cultures on micropatterned PDMS (µPs) (*Figure 1, Figure 2A*). Prior to protein extraction, hiPSC-CFs were selectively depleted from samples using a method adapted from protocols for the selective adhesion of primary fibroblasts from tissue (*Supplemental Methods*).^31^ Briefly, solubilized samples were incubated on uncoated TCP wells for one hour. Due to their limited adherence capacity, the samples’ cardiomyocytes cannot attach to uncoated TCP in such short time periods,^32^ they remained in suspension while the fibroblasts adhered to the TCP surface. This process yielded enriched hiPSC-CM samples of approximately 50,000 cells to be treated with the optimized protocol. On the day of LC-MS analysis, proteins were extracted from the cell pellets by the addition of 20 µL of the 90% HFIP extraction solution, generating complex, whole-cell lysates.

**FIGURE 1.**
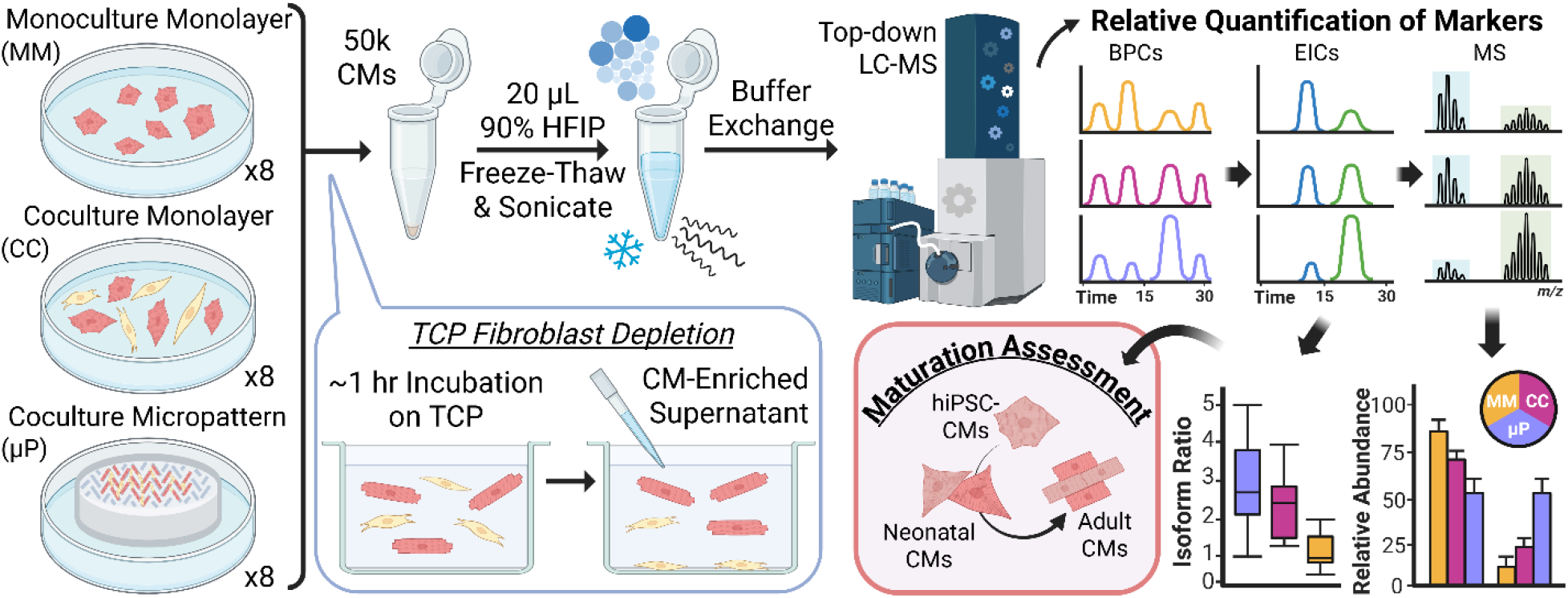
Experimental overview for the maturation assessment of micropatterned hiPSC-CMs in comparison to hiPSC-CMs cultured on non-patterned substrate. Three groups (n=8 each) of human induced pluripotent stem cell-derived cardiomyocytes (hiPSC-CMs) were cultured for analysis: monoculture CMs on non-patterned substrate (MM), CMs cocultured alongside hiPSC-derived cardiac fibribroblasts (CFs) on non-patterned substrate (CC), and CMs cocultured with CFs on the micropatterned substrate (µP). After depletion of CFs using their unique ability to quickly adhere to uncoared tissue culture plastic (TCP) plates, sarcomeric proteins were extracted from just 50,000 enriched CMs using an optimized surfactant-free protocol, followed immediately by top-down proteomic analysis.

**FIGURE 2.**
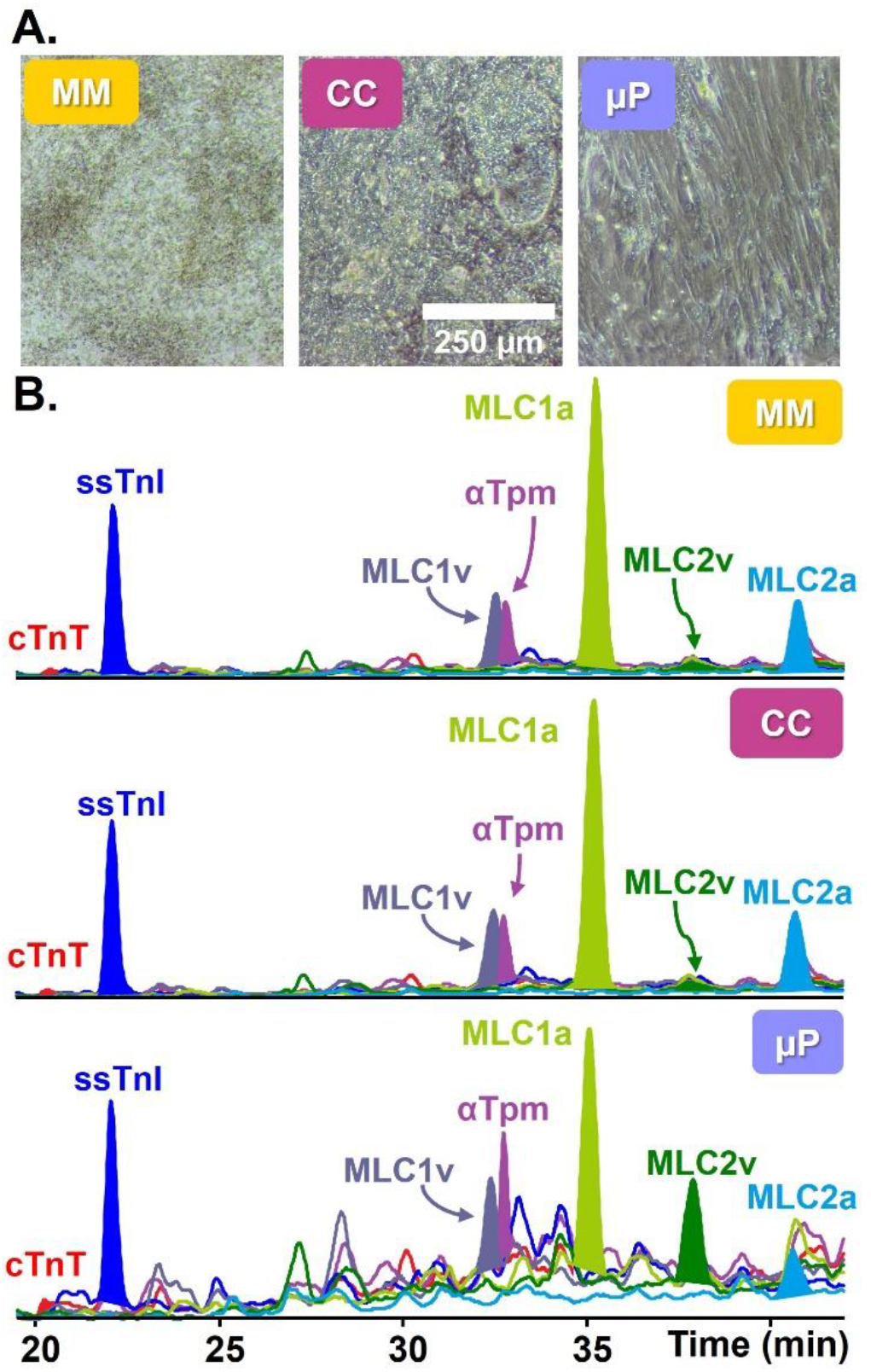
Assessment of changes in isoform expression associated with maturation. **(A)** Light microscope (10X) images of Day 14 MM, CC, and µP cultures showing the organized growth of µP hiPSC-CMs enabling elongated, rod-like cellular shapes more consistent with adult cardiomyocyte phenotypes. **(B)** Representative extraction ion chromatograms from MM, CC, µP cohorts showing consistent sarcomere protein elution times along with changes in relative expression between atrial and ventricular isoforms of myosin light chain.

From this complex mixture, we successfully identified and quantified key sarcomeric proteins in each sample at the MS1 level: the canonical adult isoform of troponin T (cTnT6, *TNNT2*), slow-skeletal troponin I (ssTnI, *TNNI1*), the ventricular isoform of MLC 1 (MLC1v, *MYL3*), alpha-tropomyosin (α-Tpm, *TPM1*), the atrial isoform of MLC 1 (MLC1a, *MYL4*), the ventricular isoform of MLC 2 (MLC2v, *MYL2*), the atrial isoform of MLC 2 (MLC2a, *MYL7*), and troponin C (TnC, *TNNC1*) (*Supplementary Figure S3*). The abundance of each protein is represented by the area under the curve (AUC) of their extracted ion chromatogram (EIC), and this area was used to compare the relative abundance of protein isoforms as ratios for cross-sample and cross-sample type comparison. Representative EICs of each group identified consistent inter-group elution times of sarcomere proteins while also highlighting notable changes in relative MLC isoform abundance between groups (*Figure 2B*). Additionally, the base peak chromatograms (BPCs), or a trace of the most abundant ion eluting from the column as a function of time, of the MM, CC, and µP groups (*Supplementary Figures S4-S6*) reveal consistent intra-group elution times, confirming reproducible extraction efficiency across samples and enabling reliable relative quantification within this study.

### Relative quantification of changes in isoform expression associated with maturation

Our previous top-down proteomics studies have demonstrated that cardiomyocyte maturation is marked by specific sarcomeric protein isoform switches.^10,25^ Notably, we have previously found that dephosphorylation of α-Tpm and isoform switching from fetal to adult troponin isoforms such as the transition from ssTnI to ctnI as well as cTnT’s fetal isoform (cTnT1) to the adult cTnT6. Additionally, there is a shift from atrial to ventricular MLC isoforms, specifically the transition of MLC1a and MLC2a to MLC1v and MLC2v, respectively.^10,25^ However, our data revealed that the MLCv/MLC1a ratio did not exhibit a significant change between MM and CC samples. By contrast, a significant increase was observed in the µPs when compared to both MMs and CCs (*Supplementary Figure S7)*. Similarly, the ratio of MLC2v/MLC2a was significantly elevated in µPs whereas no statistically significant differences were observed when compared to MMs and CCs (*Figure 3*). We note that higher variability was detected in the µPs compared to MMs and CCs, likely due to the increase in model complexity inherently introducing more parameters for inter-sample variation.

**FIGURE 3.**
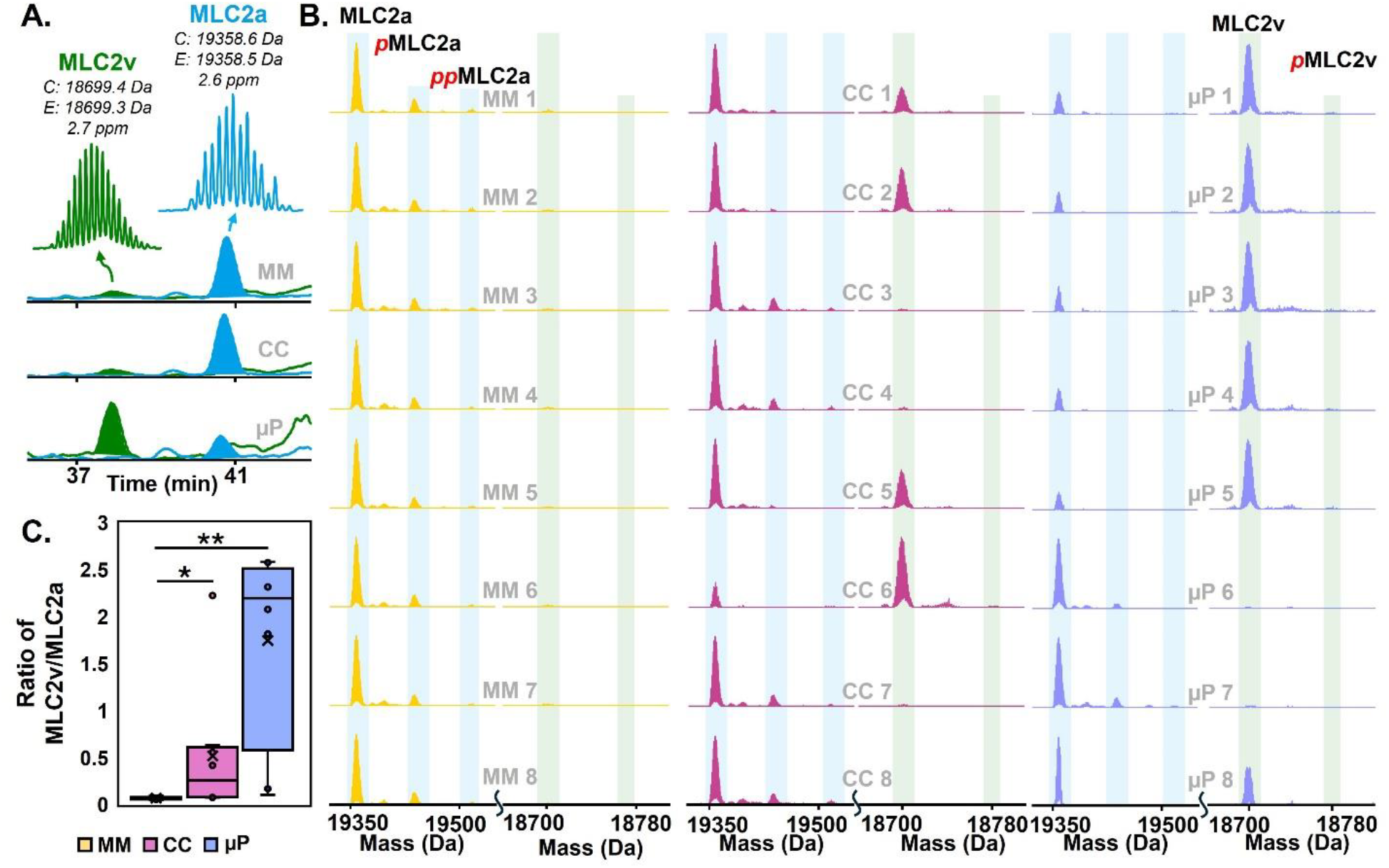
Top-down LC-MS quantitative analysis of myosin light chain 2 ventricular and atrial isoforms. A) Representative EICs of MM, CC, and µP cohorts displaying the relative abundance of MLC2a and MLC2v. The shaded portion indicates the AUC used for quantitation of isoform ratio. Above each EIC, a color-matched representation of the isotopic resolution of each isoform with the calculated and experimental most abundant masses. B) Deconvoluted spectra of MLC2a and MLC2v in each sample across cohorts. C) Box plot displaying the ratio of MLC2v to MLC2a, where “n.s.” indicates a *p* > 0.05, * indicates *p* < 0.05, and ** indicates a *p* < 0.001.

In addition to the MLC isoform switching, we observed changes to the expression of cTnT isoforms across cohorts. The fetal isoform cTnT1 was absent from µPs while it was robustly expressed in MMs and CCs (*Supplementary Figure S8*). The adult isoform of troponin T (cTnT6) was consistently expressed across all sample groups in its monophosphorylated form. Interestingly, we observed that the ratio of adult isoforms changed between culture treatments. In µPs, we identified a significant increase in the relative expression of cTnT6 compared to the non-canonical troponin T isoform 11 (cTnT11) compared to both MMs and CCs. In contrast, we did not observe significant differences in the cTnT6/cTnT11 ratio between the MM and CC groups. Though the ratio of these isoforms has not previously been explored as a marker of maturation in hiPSC-CMs, the absence of cTnT11 in non-failing adult left ventricular tissue and the notably larger ratio of cTnT6/cTnT11 peaks in mass spectra (non-quantified) taken from late-stage (76 days) ECTs^10^ might suggest an unexplored association with maturation.

Lastly, adult isoform cTnI was not quantifiable in any samples, though its monophosphorylated proteoform was sporadically detected at low abundances (peak signal intensity <6000) exclusively in µPs (*Supplementary Figure S9*). We did not detect cTnI in MMs or CCs at any point during the data collection process. The lack of reproducible cTnI identification in µPs could indicate that variation in the molecular maturity of µPs results in insufficiently reproducible cTnI expression for analysis. Alternatively, this could suggest that the limit of detection of the instrument is insufficient to account for such a low abundance signal in the complex µP lysates. While the absence of quantifiable cTnI in the µPs might suggest a less mature model, we have previously established that cTnI expression typically increases after prolonged post-differentiation culture durations, which is consistent with the observed low abundance observed in this study at Day 14 post-differentiation.^10,23^

### Quantitative analysis of the relative abundance of sarcomeric protein PTMs by top-down proteomics

In addition to changes in isoform abundance, this optimized protocol enabled PTM-level quantification, which we utilized to determine the relative abundance of phosphorylation in α-Tpm, MLC2a, MLC2v, and ssTnI (*Figure 4*). In particular, we had a specific focus on the phosphorylation of α-Tpm, an identified maturation marker.^10^ No statistically significant difference in α-Tpm phosphorylation was observed between MMs and CCs; however µPs exhibited a significant reduction in phosphorylation relative to MMs and CCs (*Supplementary Figure S10)*. There were also notable changes in the relative abundance of phosphorylation in the MLC2 isoforms. For MLC2a, both mono- and bisphosphorylated proteoforms (*p*MLC2a and *pp*MLC2a, respectively) were detected. A significant difference in the relative abundance of MLC2a and *pp*MLC2a was found between µPs and MMs, whereas µPs and CCs were only differentially expressed for the *p*MLC2a. The total phosphorylation of each group reveals a significant trend with decreasing total phosphorylation in order MMs > CCs > µPs, which is consistent with previous findings that MLC2a phosphorylation decreases with increased culture time and advanced maturation.^10^ In contrast, µPs demonstrated a significant increase in MLC2v phosphorylation when compared to both MMs and CCs, with the *p*MLC2v proteoform notably absent from the MM cohort altogether. Finally, the phosphorylation of ssTnI was not differentially expressed in MMs versus CCs; however, µPs had a significant increase in the relative phosphorylation of ssTnI compared to both groups.

**FIGURE 4.**
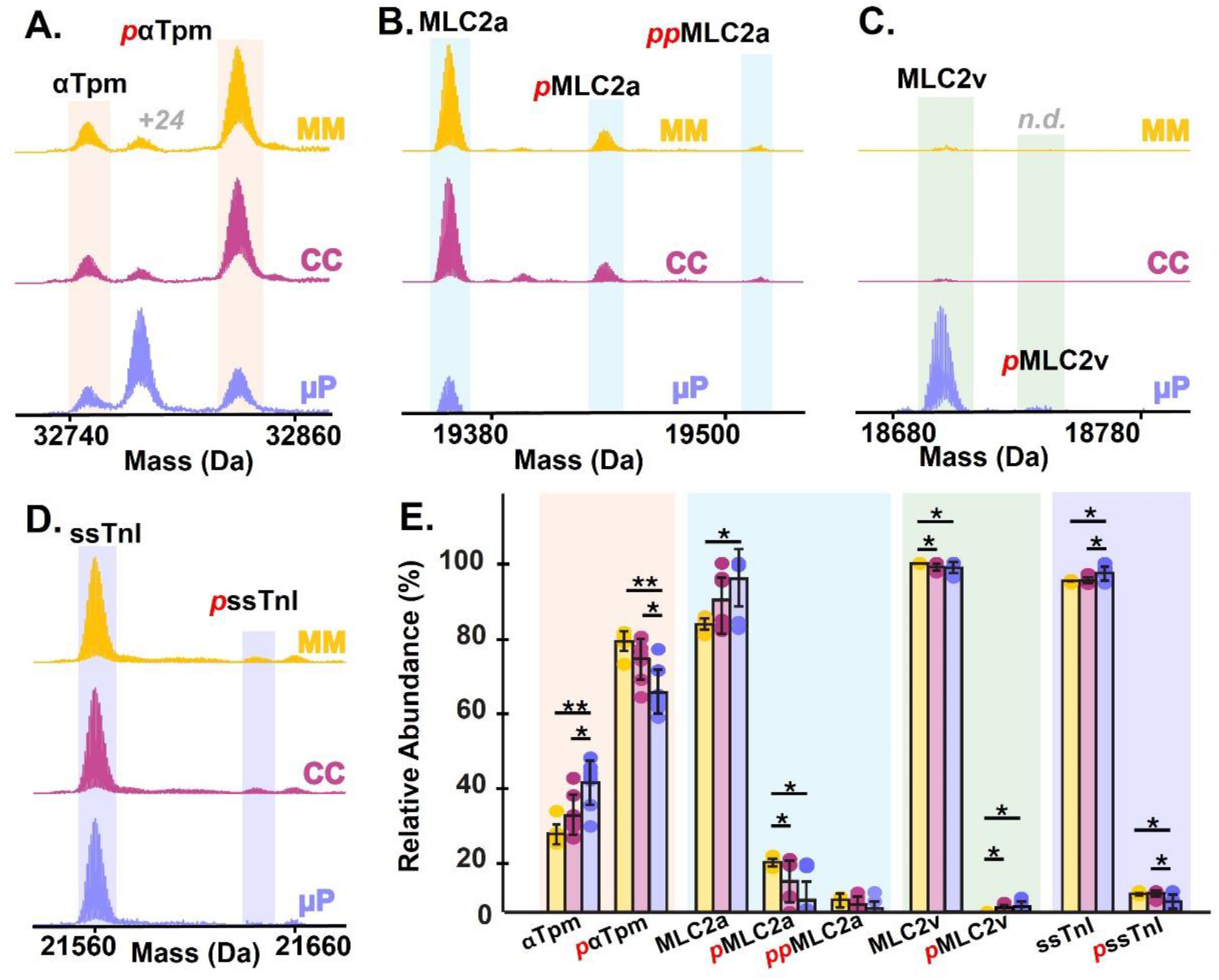
Quantitative analysis of the relative abundance of sarcomeric protein post-translational modifications by top-down proteomics. **(A-D)** Top-down mass spectra of sarcomere proteins alpha-Tropomyosin (α-Tpm), myosin light chain 2 atrial isoform (MLC2a), myosin light chain 2 ventricular isoform (MLC2v), and slow-skeletal troponin I (ssTnI). “n.d.” indicates that the proteoform was not detected. **(E)** Quantitation of proteoform relative abundance, where * indicates *p* < 0.05, and ** indicates a *p* < 0.001. The colored boxes correspond with the associated proteins highlighted in A-D.

Collectively, these findings indicate that hiPSC-CMs co-cultured on micropatterns exhibit more mature molecular phenotypes than traditional monoculture and co-culture monolayer hiPSC-CMs and are comparable in maturity to other *in vitro* cardiac models popular for proteomic studies, such as 3D constructs. Though this study did not include a comparison of µPs with 3D constructs our prior research has reported extensively on their maturity through sarcomeric maturation markers.^24^ Previously, we have reported that 3D constructs exhibit enhanced maturation when compared with monocultured hiPSC-CMs from the same cell line. Interestingly, the relative abundance of phosphorylated α-Tpm was comparable to phosphorylation observed in both ESC-derived CMs cultured for 60 days^10^ and in 3D constructs created with the DF19-9-11T line.^24^ Further, we observed that µPs exhibited a *higher* ratio of MLC1v/MLC1a than co-cultured ECT derived from the same DF19-911T line.^23^ Additionally, previous works have indicated that co-culture alongside CFs accelerates sarcomere maturation of hiPSC-CM models.^16,23^ Despite this, our work indicates that the presence of CFs alone did not induce advanced maturation during the 14 days of co-culture on micropatterns (post-differentiation), as evident through the absence of significant differences between MMs and CCs. This supports our conclusion that the enhanced maturation observed in µPs is driven by the guided spatial organization directed by the patterned PDMS substrate.

## CONCLUSIONS

In this study, we have established a surfactant-free protein extraction method tailored for high-sensitivity top-down proteomics of limited, heterogeneous multi-cell samples. We applied this method to assess the maturation of micropatterned hiPSC-CMs and observed proteoform-level maturation signatures such as the increased expression of ventricular MLC isoforms, the decreased phosphorylation of α-Tpm, and isoform switching from fetal cTnT to adult cTnT. Interestingly, we also identified an increase in the ratio of cTnT6/cTnT11 in micropatterned hiPSC-CMs, which has not yet been characterized as a marker of sarcomere maturation, potentially implicating non-canonical cTnT isoforms in a developmental role. These proteomic findings confirm that the micropatterned hiPSC-CMs exhibit more mature proteoform and isoform expression than co-culture and monocultured hiPSC-CMs alone and provides a promising platform for use in cardiac disease and arrhythmogenic studies.

## ASSOCIATED DATA

All data of this study is available from the corresponding author upon reasonable request.

## ACKNOWLEDGEMENTS

Financial support was provided by National Institutes of Health (NIH) R01 HL170521 (to L.L.E.). Y.G. would also like to acknowledge R01 HL096971 and HL109810. M.C.W. acknowledges that this material is based upon work supported by the National Science Foundation Graduate Research Fellowship Program under grant no. 2137424. M.J. acknowledges support provided by the National Institutes of Health, under the Ruth L. Kirschstein National Research Service Award T32HL007936 from the National Heart Lung and Blood Institute to the University of Wisconsin-Madison Cardiovascular Research Center. K.J.R. acknowledges the National Science Foundation Graduate Research Fellowship Program under grant no. DGE-1747503 and the Graduate School and the Office of the Vice Chancellor for Research and Graduate Education at the University of Wisconsin-Madison, funded by Wisconsin Alumni Research Foundation. Any opinions, findings, and conclusions or recommendations expressed in this material are those of the author(s) and do not necessarily reflect the views of the National Science Foundation. The authors would like to acknowledge the UW-Madison Human Proteomics Program Mass Spectrometry Facility (initially funded by the Wisconsin partnership funds) for support in obtaining MS data and NIH S10OD018475 for the acquisition of the ultra–high-resolution mass spectrometer for biomedical research. All authors would like to acknowledge the hiPSC 19-9-11 cell line donor for their gracious gift to this project and the stem cell research community. We also want to thank Holden Rogers, who provided helpful discussions.

## CONFLICT OF INTEREST STATEMENT

TJK is a consultant for Fujifilm Cellular Dynamics Incorporated.

## REFERENCES

(1) Hnatiuk, A. P.; Briganti, F.; Staudt, D.; Mercola, M. Human iPSC Modeling of Heart Disease for Drug Development. Cell Chem Biol 2021, 28 (3), 271–282. 10.1016/j.chembiol.2021.02.016.

(2) Zhang, J.; Tao, R.; Campbell, K. F.; Carvalho, J. L.; Ruiz, E. C.; Kim, G. C.; Schmuck, E. G.; Raval, A. N.; da Rocha, A. M.; Herron, T. J.; Jalife, J.; Thomson, J. A.; Kamp, T. J. Functional Cardiac Fibroblasts Derived from Human Pluripotent Stem Cells via Second Heart Field Progenitors. Nat Commun 2019, 10 (1), 2238. 10.1038/s41467-019-09831-5.

(3) Matsa, E.; Burridge, P. W.; Wu, J. C. Human Stem Cells for Modeling Heart Disease and for Drug Discovery. Science Translational Medicine 2014, 6 (239), 239ps6–239ps6. 10.1126/scitranslmed.3008921.

(4) van Meer, B. J.; Tertoolen, L. G. J.; Mummery, C. L. Concise Review: Measuring Physiological Responses of Human Pluripotent Stem Cell Derived Cardiomyocytes to Drugs and Disease. Stem Cells 2016, 34 (8), 2008–2015. 10.1002/stem.2403.

(5) Lalit, P. A.; Hei, D. J.; Raval, A. N.; Kamp, T. J. Induced Pluripotent Stem Cells for Post-Myocardial Infarction Repair: Remarkable Opportunities and Challenges. Circ Res 2014, 114 (8), 1328–1345. 10.1161/CIRCRESAHA.114.300556.

(6) Chen, I. Y.; Matsa, E.; Wu, J. C. Induced Pluripotent Stem Cells: At the Heart of Cardiovascular Precision Medicine. Nat Rev Cardiol 2016, 13 (6), 333–349. 10.1038/nrcardio.2016.36.

(7) Jebran, A.-F.; Seidler, T.; Tiburcy, M.; Daskalaki, M.; Kutschka, I.; Fujita, B.; Ensminger, S.; Bremmer, F.; Moussavi, A.; Yang, H.; Qin, X.; Mißbach, S.; Drummer, C.; Baraki, H.; Boretius, S.; Hasenauer, C.; Nette, T.; Kowallick, J.; Ritter, C. O.; Lotz, J.; Didié, M.; Mietsch, M.; Meyer, T.; Kensah, G.; Krüger, D.; Sakib, M. S.; Kaurani, L.; Fischer, A.; Dressel, R.; Rodriguez-Polo, I.; Stauske, M.; Diecke, S.; Maetz-Rensing, K.; Gruber-Dujardin, E.; Bleyer, M.; Petersen, B.; Roos, C.; Zhang, L.; Walter, L.; Kaulfuß, S.; Yigit, G.; Wollnik, B.; Levent, E.; Roshani, B.; Stahl-Henning, C.; Ströbel, P.; Legler, T.; Riggert, J.; Hellenkamp, K.; Voigt, J.-U.; Hasenfuß, G.; Hinkel, R.; Wu, J. C.; Behr, R.; Zimmermann, W.-H. Engineered Heart Muscle Allografts for Heart Repair in Primates and Humans. Nature 2025, 639 (8054), 503–511. 10.1038/s41586-024-08463-0.

(8) Ang, Y.-S.; Rivas, R. N.; Ribeiro, A. J. S.; Srivas, R.; Rivera, J.; Stone, N. R.; Pratt, K.; Mohamed, T. M. A.; Fu, J.-D.; Spencer, C. I.; Tippens, N. D.; Li, M.; Narasimha, A.; Radzinsky, E.; Moon-Grady, A. J.; Yu, H.; Pruitt, B. L.; Snyder, M. P.; Srivastava, D. Disease Model of GATA4 Mutation Reveals Transcription Factor Cooperativity in Human Cardiogenesis. Cell 2016, 167 (7), 1734-1749.e22. 10.1016/j.cell.2016.11.033.

(9) Sayed, N.; Liu, C.; Wu, J. C. Translation of Human-Induced Pluripotent Stem Cells: From Clinical Trial in a Dish to Precision Medicine. Journal of the American College of Cardiology 2016, 67 (18), 2161–2176. 10.1016/j.jacc.2016.01.083.

(10) Cai, W.; Zhang, J.; de Lange, W. J.; Gregorich, Z. R.; Karp, H.; Farrell, E. T.; Mitchell, S. D.; Tucholski, T.; Lin, Z.; Biermann, M.; McIlwain, S. J.; Ralphe, J. C.; Kamp, T. J.; Ge, Y. An Unbiased Proteomics Method to Assess the Maturation of Human Pluripotent Stem Cell–Derived Cardiomyocytes. Circulation Research 2019, 125 (11), 936–953. 10.1161/CIRCRESAHA.119.315305.

(11) Peters, N. S.; Severs, N. J.; Rothery, S. M.; Lincoln, C.; Yacoub, M. H.; Green, C. R. Spatiotemporal Relation between Gap Junctions and Fascia Adherens Junctions during Postnatal Development of Human Ventricular Myocardium. Circulation 1994, 90 (2), 713–725. 10.1161/01.CIR.90.2.713.

(12) Nunes, S. S.; Miklas, J. W.; Liu, J.; Aschar-Sobbi, R.; Xiao, Y.; Zhang, B.; Jiang, J.; Massé, S.; Gagliardi, M.; Hsieh, A.; Thavandiran, N.; Laflamme, M. A.; Nanthakumar, K.; Gross, G. J.; Backx, P. H.; Keller, G.; Radisic, M. Biowire: A Platform for Maturation of Human Pluripotent Stem Cell–Derived Cardiomyocytes. Nat Methods 2013, 10 (8), 781–787. 10.1038/nmeth.2524.

(13) Shadrin, I. Y.; Allen, B. W.; Qian, Y.; Jackman, C. P.; Carlson, A. L.; Juhas, M. E.; Bursac, N. Cardiopatch Platform Enables Maturation and Scale-up of Human Pluripotent Stem Cell-Derived Engineered Heart Tissues. Nat Commun 2017, 8 (1), 1825. 10.1038/s41467-017-01946-x.

(14) Li, W.; Luo, X.; Strano, A.; Arun, S.; Gamm, O.; Poetsch, M. S.; Hasse, M.; Steiner, R.-P.; Fischer, K.; Pöche, J.; Ulbricht, Y.; Lesche, M.; Trimaglio, G.; El-Armouche, A.; Dahl, A.; Mirtschink, P.; Guan, K.; Schubert, M. Comprehensive Promotion of iPSC-CM Maturation by Integrating Metabolic Medium with Nanopatterning and Electrostimulation. Nat Commun 2025, 16 (1), 2785. 10.1038/s41467-025-58044-6.

(15) Yang, H.; Yang, Y.; Kiskin, F. N.; Shen, M.; Zhang, J. Z. Recent Advances in Regulating the Proliferation or Maturation of Human-Induced Pluripotent Stem Cell-Derived Cardiomyocytes. Stem Cell Res Ther 2023, 14, 228. 10.1186/s13287-023-03470-w.

(16) Reilly, L.; Munawar, S.; Zhang, J.; Crone, W. C.; Eckhardt, L. L. Challenges and Innovation: Disease Modeling Using Human-Induced Pluripotent Stem Cell-Derived Cardiomyocytes. Front. Cardiovasc. Med. 2022, 9. 10.3389/fcvm.2022.966094.

(17) Napiwocki, B. N.; Stempien, A.; Lang, D.; Kruepke, R. A.; Kim, G.; Zhang, J.; Eckhardt, L. L.; Glukhov, A. V.; Kamp, T. J.; Crone, W. C. Micropattern Platform Promotes Extracellular Matrix Remodeling by Human PSC-Derived Cardiac Fibroblasts and Enhances Contractility of Co-Cultured Cardiomyocytes. Physiol Rep 2021, 9 (19), e15045. 10.14814/phy2.15045.

(18) Stempien, A.; Josvai, M.; de Lange, W. J.; Hernandez, J. J.; Notbohm, J.; Kamp, T. J.; Valdivia, H. H.; Eckhardt, L. L.; Maginot, K. R.; Ralphe, J. C.; Crone, W. C. Identifying Features of Cardiac Disease Phenotypes Based on Mechanical Function in a Catecholaminergic Polymorphic Ventricular Tachycardia Model. Front Bioeng Biotechnol 2022, 10, 873531. 10.3389/fbioe.2022.873531.

(19) Aebersold, R.; Agar, J. N.; Amster, I. J.; Baker, M. S.; Bertozzi, C. R.; Boja, E. S.; Costello, C. E.; Cravatt, B. F.; Fenselau, C.; Garcia, B. A.; Ge, Y.; Gunawardena, J.; Hendrickson, R. C.; Hergenrother, P. J.; Huber, C. G.; Ivanov, A. R.; Jensen, O. N.; Jewett, M. C.; Kelleher, N. L.; Kiessling, L. L.; Krogan, N. J.; Larsen, M. R.; Loo, J. A.; Ogorzalek Loo, R. R.; Lundberg, E.; MacCoss, M. J.; Mallick, P.; Mootha, V. K.; Mrksich, M.; Muir, T. W.; Patrie, S. M.; Pesavento, J. J.; Pitteri, S. J.; Rodriguez, H.; Saghatelian, A.; Sandoval, W.; Schlüter, H.; Sechi, S.; Slavoff, S. A.; Smith, L. M.; Snyder, M. P.; Thomas, P. M.; Uhlén, M.; Van Eyk, J. E.; Vidal, M.; Walt, D. R.; White, F. M.; Williams, E. R.; Wohlschlager, T.; Wysocki, V. H.; Yates, N. A.; Young, N. L.; Zhang, B. How Many Human Proteoforms Are There? Nat Chem Biol 2018, 14 (3), 206–214. 10.1038/nchembio.2576.

(20) Smith, L. M.; Agar, J. N.; Chamot-Rooke, J.; Danis, P. O.; Ge, Y.; Loo, J. A.; Paša-Tolić, L.; Tsybin, Y. O.; Kelleher, N. L.; The Consortium for Top-Down Proteomics. The Human Proteoform Project: Defining the Human Proteome. Science Advances 2021, 7 (46), eabk0734. 10.1126/sciadv.abk0734.

(21) Smith, L. M.; Kelleher, N. L. Proteoform: A Single Term Describing Protein Complexity. Nat Methods 2013, 10 (3), 186–187. 10.1038/nmeth.2369.

(22) Roberts, D. S.; Loo, J. A.; Tsybin, Y. O.; Liu, X.; Wu, S.; Chamot-Rooke, J.; Agar, J. N.; Paša-Tolić, L.; Smith, L. M.; Ge, Y. Top-down Proteomics. Nat Rev Methods Primers 2024, 4 (1), 1–23. 10.1038/s43586-024-00318-2.

(23) Melby, J. A.; Roberts, D. S.; Larson, E. J.; Brown, K. A.; Bayne, E. F.; Jin, S.; Ge, Y. Novel Strategies to Address the Challenges in Top-Down Proteomics. J. Am. Soc. Mass Spectrom. 2021, 32 (6), 1278–1294. 10.1021/jasms.1c00099.

(24) Melby, J. A.; de Lange, W. J.; Zhang, J.; Roberts, D. S.; Mitchell, S. D.; Tucholski, T.; Kim, G.; Kyrvasilis, A.; McIlwain, S. J.; Kamp, T. J.; Ralphe, J. C.; Ge, Y. Functionally Integrated Top-Down Proteomics for Standardized Assessment of Human Induced Pluripotent Stem Cell-Derived Engineered Cardiac Tissues. J Proteome Res 2021, 20 (2), 1424–1433. 10.1021/acs.jproteome.0c00830.

(25) Rossler, K. J.; Lange, W. J. de; Mann, M. W.; Aballo, T. J.; Melby, J. A.; Zhang, J.; Kim, G.; Bayne, E. F.; Zhu, Y.; Farrell, E. T.; Kamp, T. J.; Ralphe, J. C.; Ge, Y. Lactate- and Immunomagnetic-Purified hiPSC–Derived Cardiomyocytes Generate Comparable Engineered Cardiac Tissue Constructs. JCI Insight 2024, 9 (1). 10.1172/jci.insight.172168.

(26) Bedada, F. B.; Chan, S. S.-K.; Metzger, S. K.; Zhang, L.; Zhang, J.; Garry, D. J.; Kamp, T. J.; Kyba, M.; Metzger, J. M. Acquisition of a Quantitative, Stoichiometrically Conserved Ratiometric Marker of Maturation Status in Stem Cell-Derived Cardiac Myocytes. Stem Cell Reports 2014, 3 (4), 594–605. 10.1016/j.stemcr.2014.07.012.

(27) Lian, X.; Zhang, J.; Azarin, S. M.; Zhu, K.; Hazeltine, L. B.; Bao, X.; Hsiao, C.; Kamp, T. J.; Palecek, S. P. Directed Cardiomyocyte Differentiation from Human Pluripotent Stem Cells by Modulating Wnt/β-Catenin Signaling under Fully Defined Conditions. Nat Protoc 2013, 8 (1), 162–175. 10.1038/nprot.2012.150.

(28) Melby, J. A.; Brown, K. A.; Gregorich, Z. R.; Roberts, D. S.; Chapman, E. A.; Ehlers, L. E.; Gao, Z.; Larson, E. J.; Jin, Y.; Lopez, J. R.; Hartung, J.; Zhu, Y.; McIlwain, S. J.; Wang, D.; Guo, W.; Diffee, G. M.; Ge, Y. High Sensitivity Top–down Proteomics Captures Single Muscle Cell Heterogeneity in Large Proteoforms. Proceedings of the National Academy of Sciences 2023, 120 (19), e2222081120. 10.1073/pnas.2222081120.

(29) Lin, Z.; Wei, L.; Cai, W.; Zhu, Y.; Tucholski, T.; Mitchell, S. D.; Guo, W.; Ford, S. P.; Diffee, G. M.; Ge, Y. Simultaneous Quantification of Protein Expression and Modifications by Top-down Targeted Proteomics: A Case of the Sarcomeric Subproteome. Mol Cell Proteomics 2019, 18 (3), 594–605. 10.1074/mcp.TIR118.001086.

(30) Yoshida, K.; Yamaguchi, T.; Adachi, T.; Otomo, T.; Matsuo, D.; Takamuku, T.; Nishi, N. Structure and Dynamics of Hexafluoroisopropanol-Water Mixtures by x-Ray Diffraction, Small-Angle Neutron Scattering, NMR Spectroscopy, and Mass Spectrometry. The Journal of Chemical Physics 2003, 119 (12), 6132–6142. 10.1063/1.1602070.

(31) Nielsen, M. S.; van Opbergen, C. J. M.; van Veen, T. A. B.; Delmar, M. The Intercalated Disc: A Unique Organelle for Electromechanical Synchrony in Cardiomyocytes. Physiological Reviews 2023, 103 (3), 2271–2319. 10.1152/physrev.00021.2022.

(32) Kumar, S.; Nagesh, D.; Ramasubbu, V.; Prabhashankar, A. B.; Sundaresan, N. R. Isolation and Culture of Primary Fibroblasts from Neonatal Murine Hearts to Study Cardiac Fibrosis. Bio Protoc 2023, 13 (4), e4616. 10.21769/BioProtoc.4616.

