## Supplemental Information for "High-Sensitivity Top-Down Proteomics Reveals Enhanced Maturation of Micropatterned Induced Pluripotent Stem Cell-Derived Cardiomyocytes"

Supplementary Methods...............................................................................................................S-3

Supplementary Figures

Supplementary Figure S1. Method optimization to establish surfactant-free extraction of proteins from limited number of cells .............................................................................S-8

Supplementary Figure S2. Technical replicates and linear instrument response analysis of the mass spectrometer .....................................................................................................S-9

Supplementary Figure S3. Top-down proteomics analysis of sarcomere proteins extracted from a representative µP sample ...................................................................................S-10

Supplementary Figure S4. Reproducibility of monoculture monolayer biological replicates ........................................................................................................................S-11

Supplementary Figure S5. Reproducibility of coculture monolayer biological

replicates ........................................................................................................................S-12

Supplementary Figure S6. Reproducibility of coculture micropattern biological

replicates ........................................................................................................................S-13

Supplementary Figure S7. Top-down LC-MS quantitation of myosin light chain 1 ventricular and atrial isoforms .......................................................................................S-14

Supplementary Figure S8. Top-down LC-MS quantitation of troponin T isoforms .....S-15

Supplementary Figure S9. Identification of low abundance adult cardiac troponin I (cTnI) by top-down LC-MS .....................................................................................................S-16

Supplementary Figure S10. Top-down LC-MS quantitation of alpha-Tropomyosin phosphorylation .............................................................................................................S-17

References .................................................................................................................................S-18

***Supplementary Methods***

Chemicals and Reagents:

All solutions were prepared with HPLC-grade water (Fisher Scientific).

A. hiPSC Culture and Maintenance

The human induced pluripotent stem cell (hiPSC) line hiPSC-DF19-9-11T (WiCell, Madison, WI) was maintained and utilized for both cardiomyocyte (CM) and fibroblast (CF) differentiation. hiPSCs were thawed and plated on Matrigel (GFR, Corning, Corning, NY) coated plates (8.7 μg/cm²) in StemFlex medium (Thermo Fisher). After reaching 50-60% confluency, hiPSCs were passaged at a lower density in StemFlex medium. Multiple passages were conducted prior to CM or CF differentiation to ensure culture homogeneity. The hiPSCs used for differentiation were between passage 40 and 60 at the onset of the differentiation process.

B. CM Differentiation and Culture

hiPSCs were differentiated to hiPSC-CMs using a modified version of the small molecule GiWi method^1^ and subsequently purified via magnetic-activated cell sorting (MACS; Miltenyi Biotec). Briefly, hiPSCs were seeded on Matrigel-coated plates and cultured in mTesR1 medium (WiCell) for 5 days, until reaching 100% confluency. On day 0 of differentiation, the culture medium was replaced with RPMI 1640 (Thermo Fisher Scientific, Waltham, MA) supplemented with 2% B27 minus insulin (Thermo Fisher Scientific) and 12 μM of the GSK3 inhibitor CHIR 99021 (Biogems, Westlake Village, CA). Twenty-four hours post-CHIR addition (day 1), the medium was changed to RPMI 1640 with 2% B27 minus insulin. On day 3, 72 hours after CHIR addition, the medium was again replaced with RPMI 1640 containing B27 minus insulin and 5 μM of the Wnt inhibitor IWP2 (StemGent, Beltsville, MD). On days 5 and 7, the cells were maintained in RPMI 1640 with B27 minus insulin. Between days 9 and 15, the medium was refreshed every other day with RPMI 1640 supplemented with 2% B27 complete medium (Thermo Fisher Scientific). Contractile activity typically began between days 9 and 12 of differentiation. On day 15, hiPSC-CMs were cryopreserved in 90% fetal bovine serum (FBS, Invitrogen, Waltham, MA) and 10% DMSO (Sigma, St. Louis, MO).

C. CF Differentiation and Culture

hiPSCs were differentiated to hiPSC-CFs as described previously.^2^ Briefly, hiPSCs were dissociated and seeded onto Matrigel-coated 6-well plates at a density of 2 × 10⁶ cells per well in mTeSR1 medium. The cells were cultured for 5 days in mTeSR1 with daily medium changes, continuing until they reached 100% confluence. On day 0 of differentiation, the medium was replaced with 2.5 mL RPMI supplemented with 2% B27 minus insulin and 12 μM CHIR. After 24 hours, on day 1, the medium was changed to 2.5 mL RPMI with 2% B27 minus insulin. On day 2, the medium was replaced with 2.5 mL of a defined fibroblast culture medium (CFBM) containing 75 ng/mL basic fibroblast growth factor (bFGF; WiCell). The cells were then maintained in CFBM supplemented with 75 ng/mL bFGF, with medium changes every other day, and cultured until day 20. hiPSC-CFs were subsequently maintained in FibroGRO (Millipore EMD, Burlington, MA) supplemented with 2% fetal bovine serum (FBS) on tissue culture plastic. Cryopreserved hiPSC-CFs were thawed and plated onto 6-well tissue culture plates at a density of 55k CFs per well in 2 mL of FibroGRO medium with 2% FBS. The cells were cultured in FibroGRO medium supplemented with 2% FBS, with medium changes every two days. hiPSC-CFs were passaged at least once after thawing before being used in experiments. All hiPSC-CFs used in this study were harvested at passage 5.

D. Elastomer Fabrication and ECM Patterning

Compliant polydimethylsiloxane (PDMS) substrates for cell seeding were fabricated by blending Sylgard 184 and Sylgard 527 (Dow Corning Corporation, Auburn, MI) in specified ratios, as described by Palchesko et al.^3^ Sylgard 184 was prepared by mixing 10 parts of base with 1 part of curing agent and stirring for 5 minutes. Sylgard 527 was prepared by combining equal volumes of components A and B and mixing for 5 minutes. To create substrates with a modulus of 10 kPa, Sylgard 184 and Sylgard 527 were combined in a 52:1 ratio, mixed for 5 minutes, and poured into a 100 mm petri dish. The uncured PDMS mixture was subjected to a vacuum for 20 minutes to remove air bubbles, after which the dish was cured for 12 hours at 60°C. Once cured, the substrate was cut into the desired dimensions using a razor blade. Mechanical testing was previously conducted to confirm that the intended PDMS modulus was achieved with this 52:1 ratio.^4^ The compliant substrates were then affixed to 12-well plates (Corning, Corning, NY) using a drop of Sylgard 184, which was cured for 6 hours at 60°C and subjected to 10 minutes of UV exposure for sterilization prior to seeding of the monolayer or patterning followed by seeding.

Microcontact printing and soft lithography were used to generate compliant substrates with patterned ECM proteins in defined geometries.^5,6,7^ A master Si wafer (FlowJEM, Toronto, ON, Canada) was used to generate reusable PDMS stamps. Sylgard 184 PDMS was cured for 6 hours at 60 °C before being cut into individual stamps for micropatterning. The stamps were coated with Matrigel and incubated at 37 °C overnight before removing excess Matrigel. Stamps were gently dried with a nitrogen airstream to remove residual moisture without dislodging the attached proteins. A polyvinyl alcohol (PVA) film was produced by mixing 0.5 g PVA beads (Sigma) with 10 mL deionized water for 20 minutes at 100 °C. The PVA solution was poured into a 100 mm petri dish and left uncovered to dry overnight. The dried film was cut into sections with an area slightly larger than the PDMS stamps and brought in contact with the dry stamps. The patterned Matrigel was allowed to transfer to the PVA film for 1 hour at 37 °C.^8^ Transfer efficiency was increased by placing a glass slide and a 50 g weight on top of each stamp. After incubation, the PVA film was removed from the stamp and brought in contact with the PDMS substrates. Matrigel was allowed to transfer to the substrate for 20 minutes at 37 °C before being washed with 3 mL phosphate-buffered saline (PBS) to dissolve the PVA film. After 10 minutes, an additional PBS wash was performed, removing the soluble PVA film and leaving only the patterned ECM proteins on the 10 kPa substrates. Substrates were maintained briefly in PBS until cell seeding.

E. hiPSC-CM and hiPSC-CF Seeding

hiPSC-CMs were thawed and plated on Matrigel at a density of 2.5 million cells per well in EB20 medium, consisting of DMEM/F12 (Life Technologies, Carlsbad, CA), 20% fetal bovine serum (FBS), 1% non-essential amino acids (NEAA), 0.5% GlutaMax, and 7 μM 2-Mercaptoethanol (Sigma). After a 48-hour recovery period, hiPSC-CMs were dissociated and singularized using TrypLE 10X (Life Technologies) for 12 minutes, followed by centrifugation at 1000 rpm for 5 minutes. Magnetic purification was then performed using the PSC-Derived Cardiomyocyte Isolation Kit (human, Miltenyi Biotec). The hiPSC-CMs were resuspended in Miltenyi Purification Buffer (PBS with 0.5% BSA and 0.4% EDTA, pH 8.0) and incubated with 20% Miltenyi Non-CM Depletion Cocktail for 5 minutes at 4°C. After centrifugation at 1000 rpm for 5 minutes, the cells were resuspended in Miltenyi Purification Buffer containing 20% Miltenyi Anti-Biotin Microbeads and incubated for 10 minutes at 4°C. The cells were then centrifuged again and resuspended in EB20 medium.

The monolayer cohorts of purified hiPSC-CMs were seeded at 2.5 × 10⁶ cells per well in 6-well plates, while micropatterned substrates were seeded with 150,000 cells per micropattern (78.5 mm²). The day of seeding was designated as day 0 of the experimental culture. Forty-eight hours after CM seeding, hiPSC-derived fibroblasts (hiPSC-CFs) were dissociated using TrypLE Express (Life Technologies) for 2 minutes and resuspended in EB20 medium. CFs were seeded at a 10:1 ratio of cardiomyocytes to fibroblasts, corresponding to 250,000 CFs in the coculture monolayer cohort (CC) or 15,000 CFs per micropattern (samples in the µP cohort).^9^ CM-only monolayers (MM) were not seeded with hiPSC-CFs. On day 3 of culture, 24 hours after CF seeding, the medium was replaced with EB20 medium, consisting of DMEM/F12, 2% FBS, 1% NEAA, 0.5% GlutaMax, and 7 μM 2-Mercaptoethanol, and the medium was exchanged every two days throughout the culture period. Samples were then harvested after 14 days.

***Supplementary Figures***

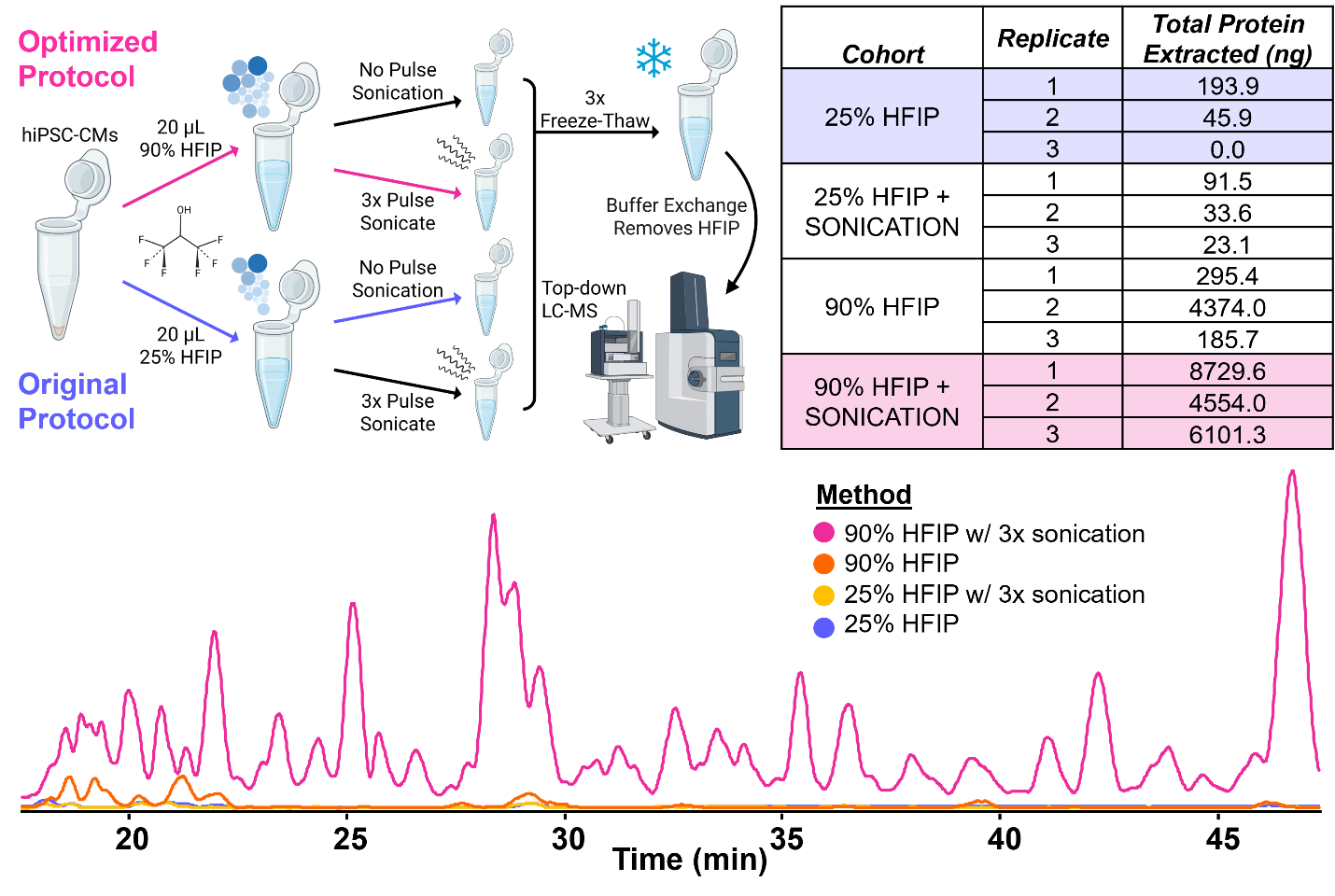

**Figure S1. Method optimization to establish surfactant-free extraction of proteins from limited number of cells.** Triplicate samples of 50k isolated hiPSC-CMs were used to identify the most effective method to extract proteins from hiPSC-CMs based on the previously developed hexafluoroisopropanol (HFIP)-enabled single muscle fiber (SMF) extraction.^10^ The optimized protocol (pink) utilizes additional homogenization and lysis via brief sonication (3x) and a higher percentage of HFIP when compared with the original protocol (purple). The table on the top right shows the total protein extracted from each sample across each cohort. Representative BPC traces from each cohort are present along the bottom of the figure. These results indicate that both protocol modifications must be added to reliably extract protein from multi-cell hiPSC-CM samples.

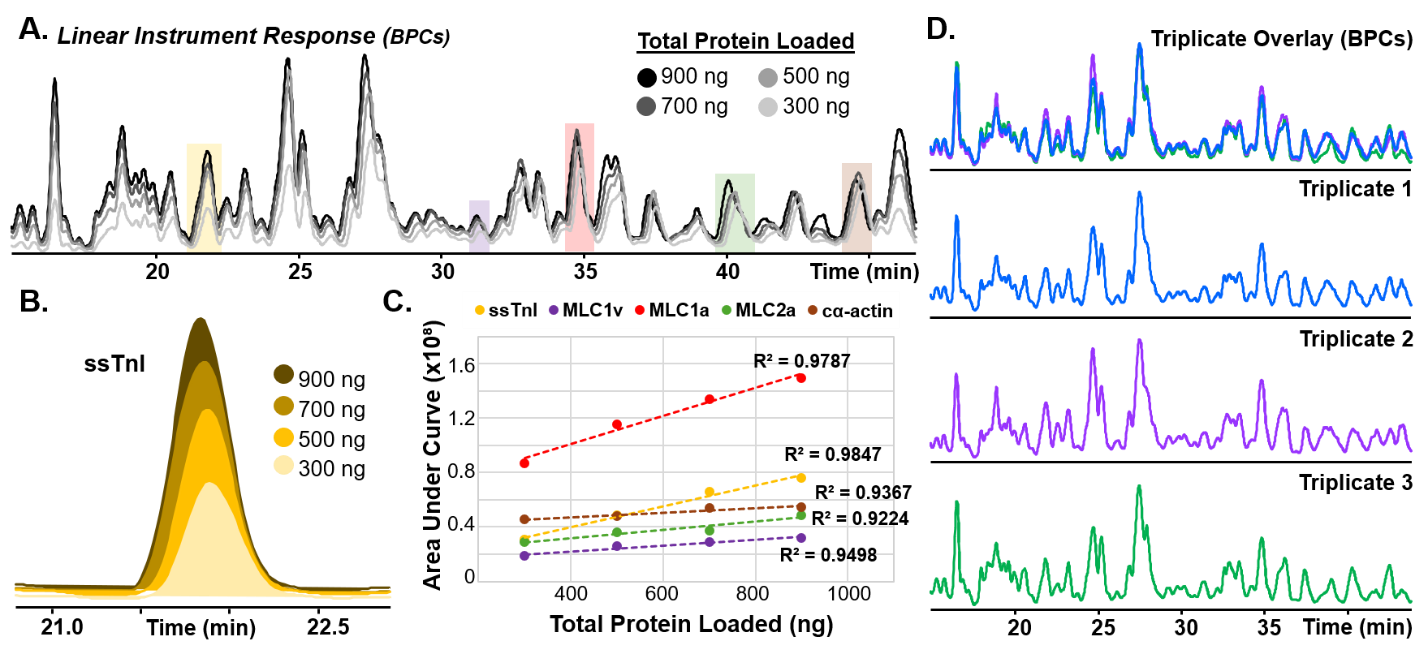

**Figure S2. High reproducibility and linear instrument response analysis of top-down LC-MS-based quantitation.** **A)** Base peak chromatograms (BPCs) of variable total protein are injected to identify the linear response range of the mass spectrometer for a hiPSC-CM sample (50k cells) using the optimized surfactant-less protocol. Elution times of select sarcomere proteins were identified the most abundant 5-7 charge state ions from the non-deconvoluted spectra were used to produce extracted ion chromatograms (EICs) across different total protein injections. **B)** EICs from slow skeletal troponin I (ssTnI) showing linearly decreasing instrument response with decreasing total protein loaded. **C)** Area under the curve of each EIC was plotted as a function of total protein loaded for each selected protein. **D)** Three technical (injection) replicates as an assessment of the stability of the mass spectrometer. The triplicate overlay of the BPCs reveals the high similarity between traces, indicating that the instrument’s response can be considered reproducible.

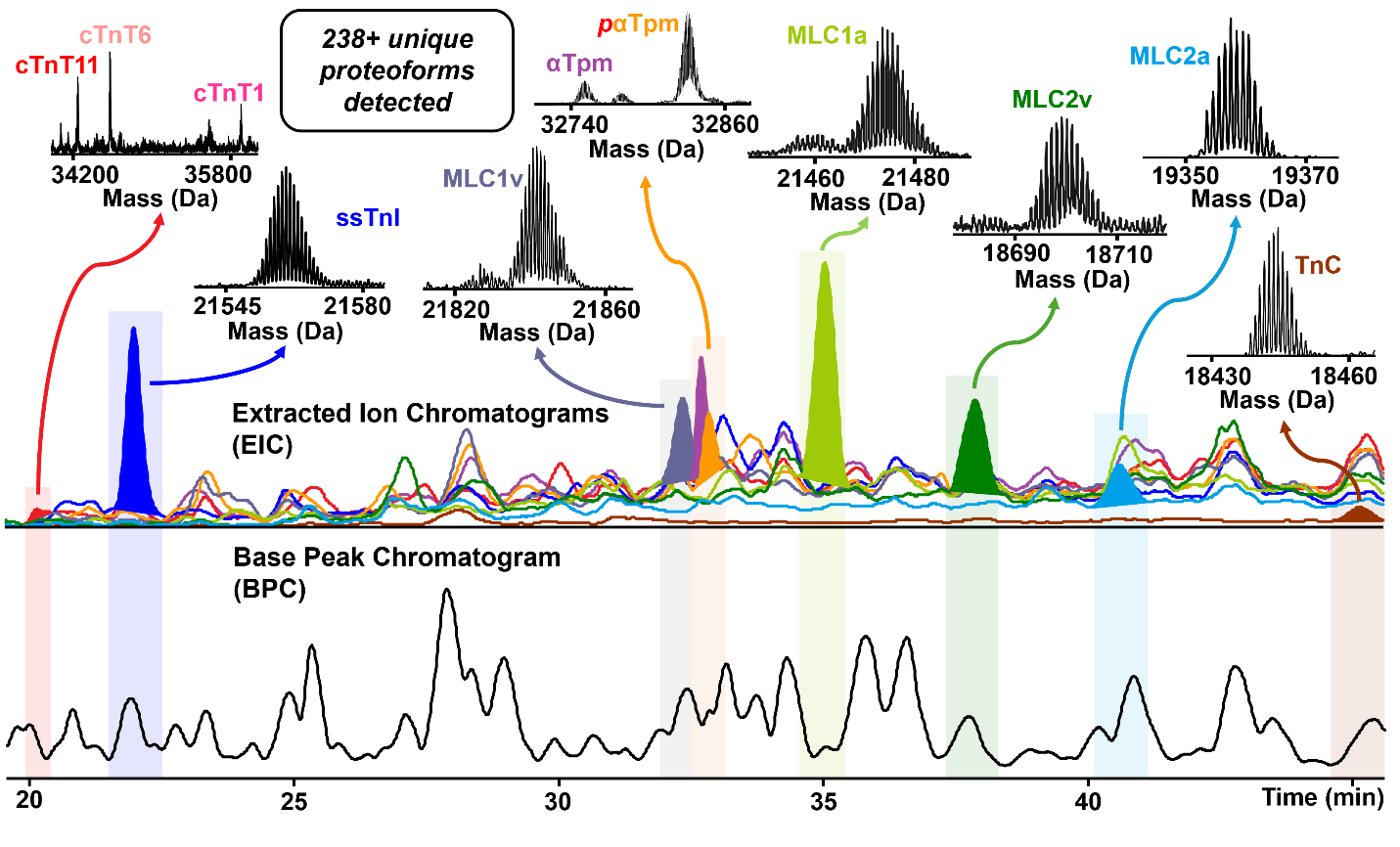

**Figure S3.** **Top-down proteomics analysis of sarcomere proteins extracted from a representative µP sample.** Base peak chromatogram (BPC) and extracted ion chromatogram (EIC) together with the deconvoluted mass spectra of sarcomere proteins are shown including troponin T (cTnT), slow-skeletal troponin I (ssTnI), myosin light chain 1 atrial and ventricular isoforms (MLC1a, MLC1v), myosin light chain 2 atrial and ventricular isoforms (MLC2a, MLC2v), alpha-Tropomyosin (α-Tpm), and troponin C (TnC).

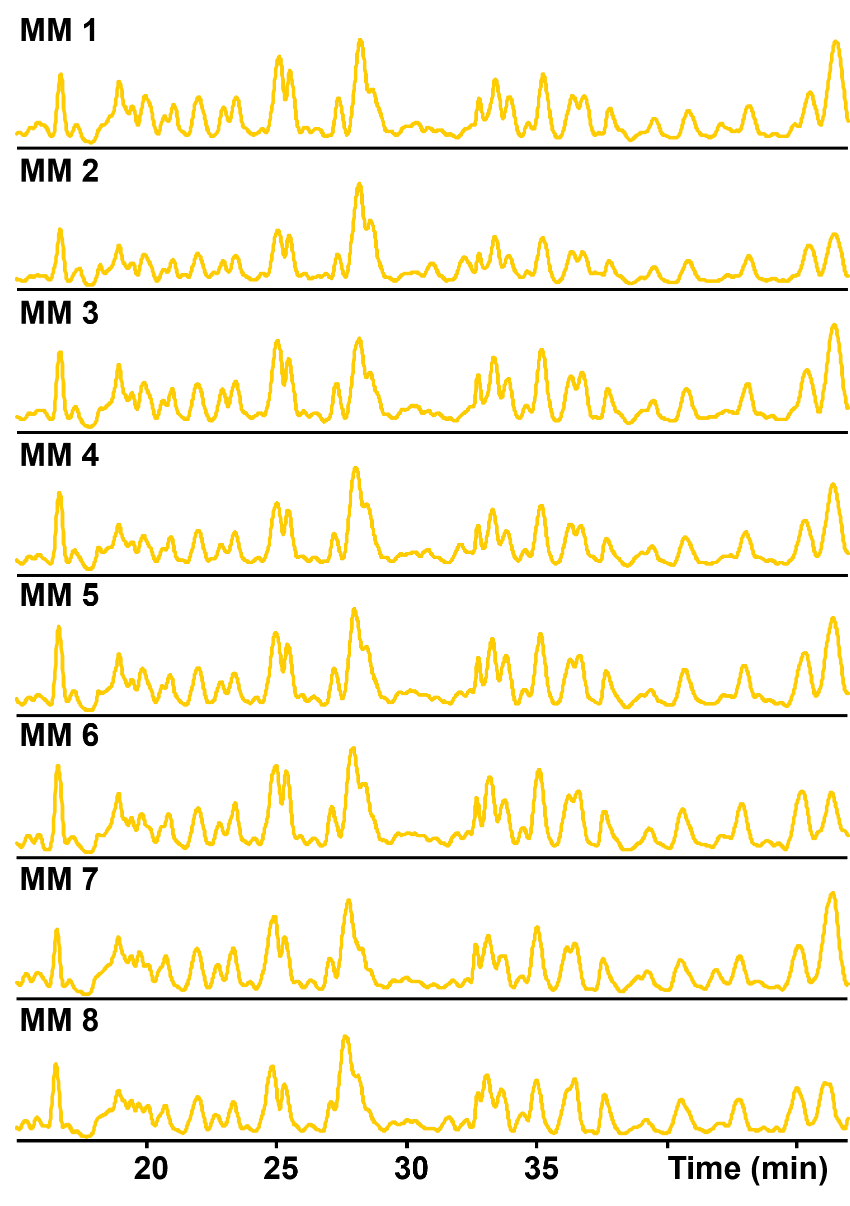

**Figure S4.** **Reproducibility of monoculture monolayer biological replicates.** BPCs of each biological replicate in the monoculture monolayer (MM) cohort (n=8) exhibit high reproducibility. Three hiPSC-CM batches are represented randomly throughout the cohort to ensure batch variability is accounted for in the analysis.

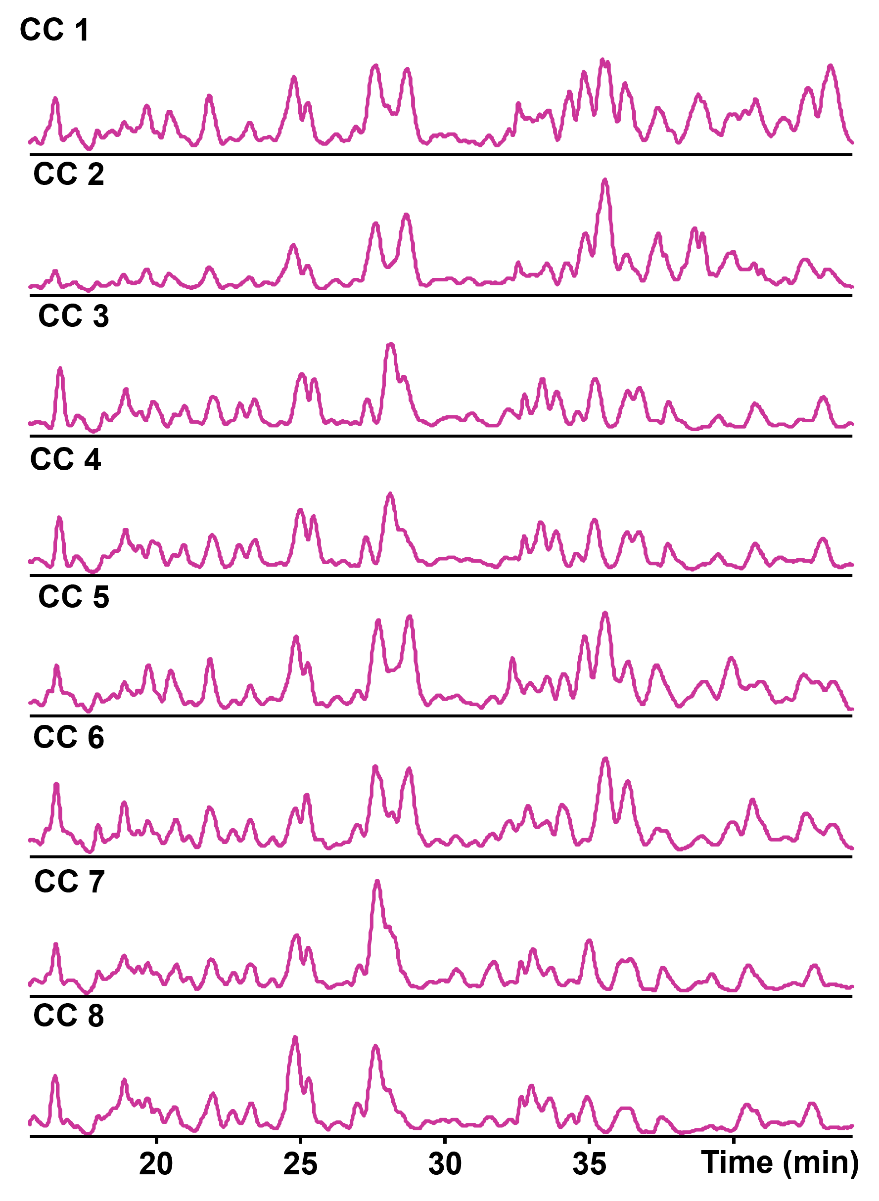

**Figure S5. Reproducibility of coculture monolayer biological replicates.** BPCs of each biological replicate in the coculture monolayer (CC) cohort (n=8) exhibit high reproducibility. Three hiPSC-CM batches are represented randomly throughout the cohort to ensure batch variability is accounted for in the analysis.

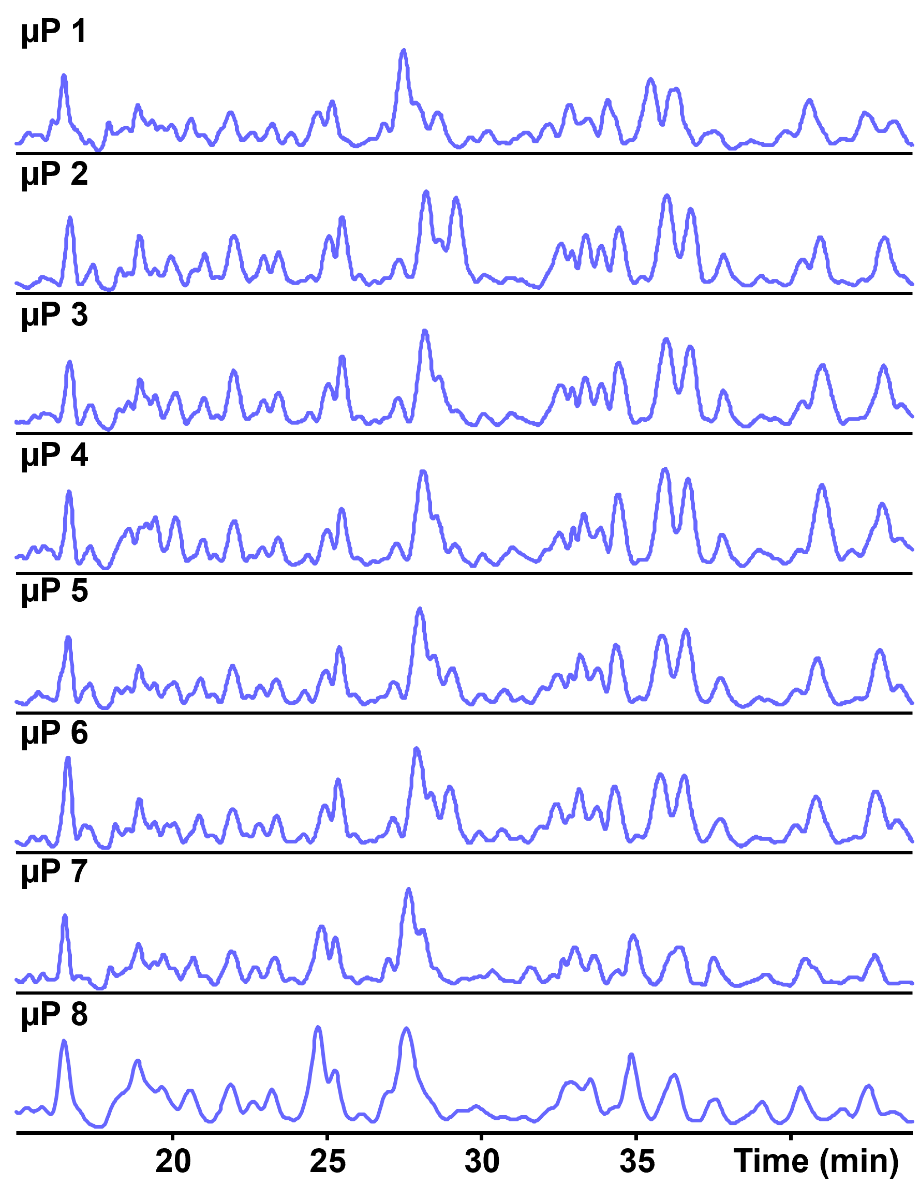

**Figure S6. Reproducibility of coculture micropattern biological replicates.** BPCs of each biological replicate in the coculture micropattern (µP) cohort (n=8) exhibit high reproducibility. Three hiPSC-CM batches are represented randomly throughout the cohort to ensure batch variability is accounted for in the analysis.

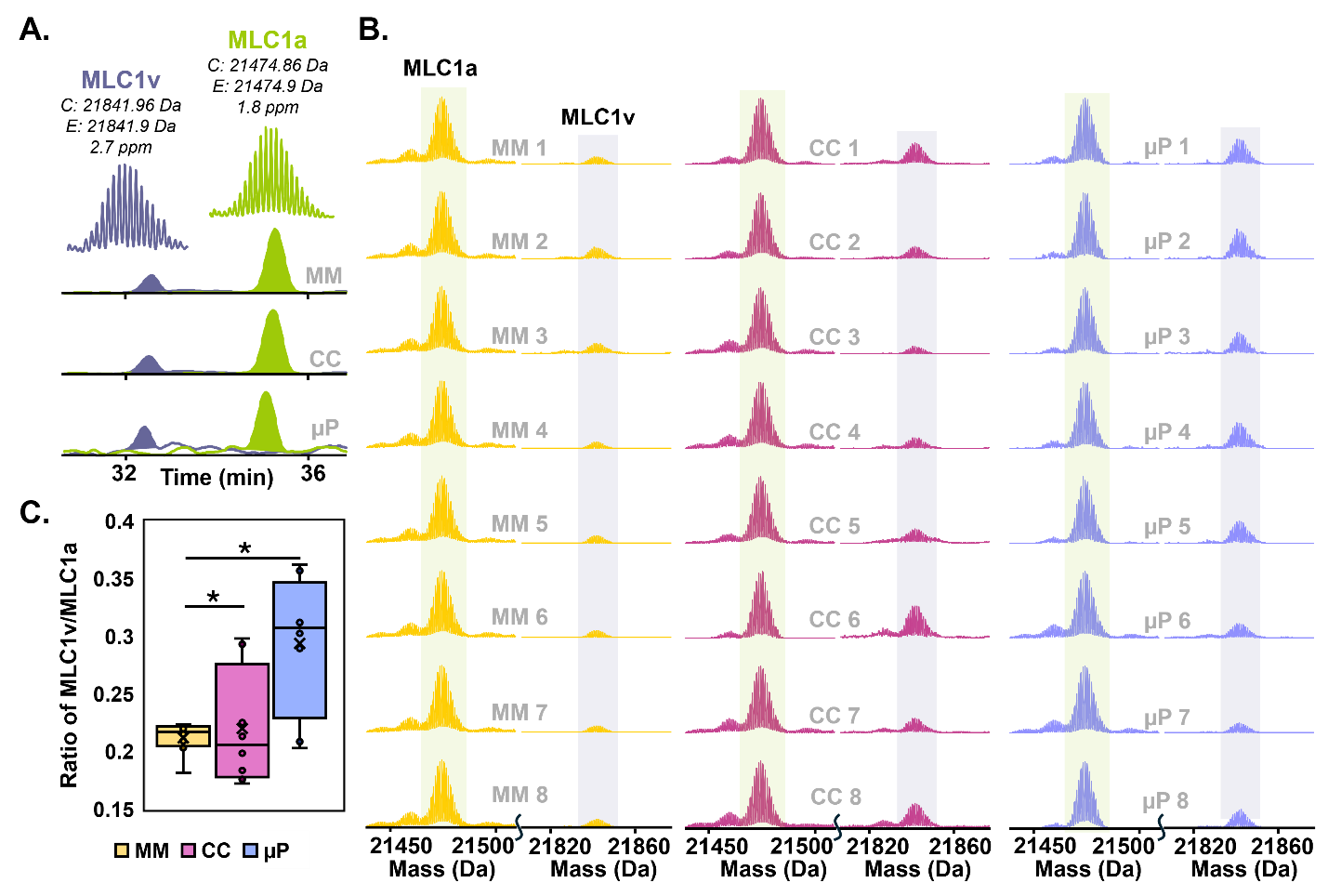

**FIGURE S7. Top-down LC-MS quantitation of myosin light chain 1 ventricular and atrial isoforms.** A) Representative EICs of MM, CC, and µP cohorts displaying the relative abundance of MLC1a and MLC1v. The shaded portion indicates the AUC used for quantitation of isoform ratio. Above each EIC, there is a color-matched representation of the isotopic resolution of each isoform with the calculated and experimental most abundant masses. B) Deconvoluted spectra of MLC1a and MLC1v in each sample across cohorts. C) Box plot displaying the ratio of MLC1v to MLC1a, where “n.s.” indicates a *p* > 0.05, * indicates *p* < 0.05, and ** indicates a *p* < 0.001.

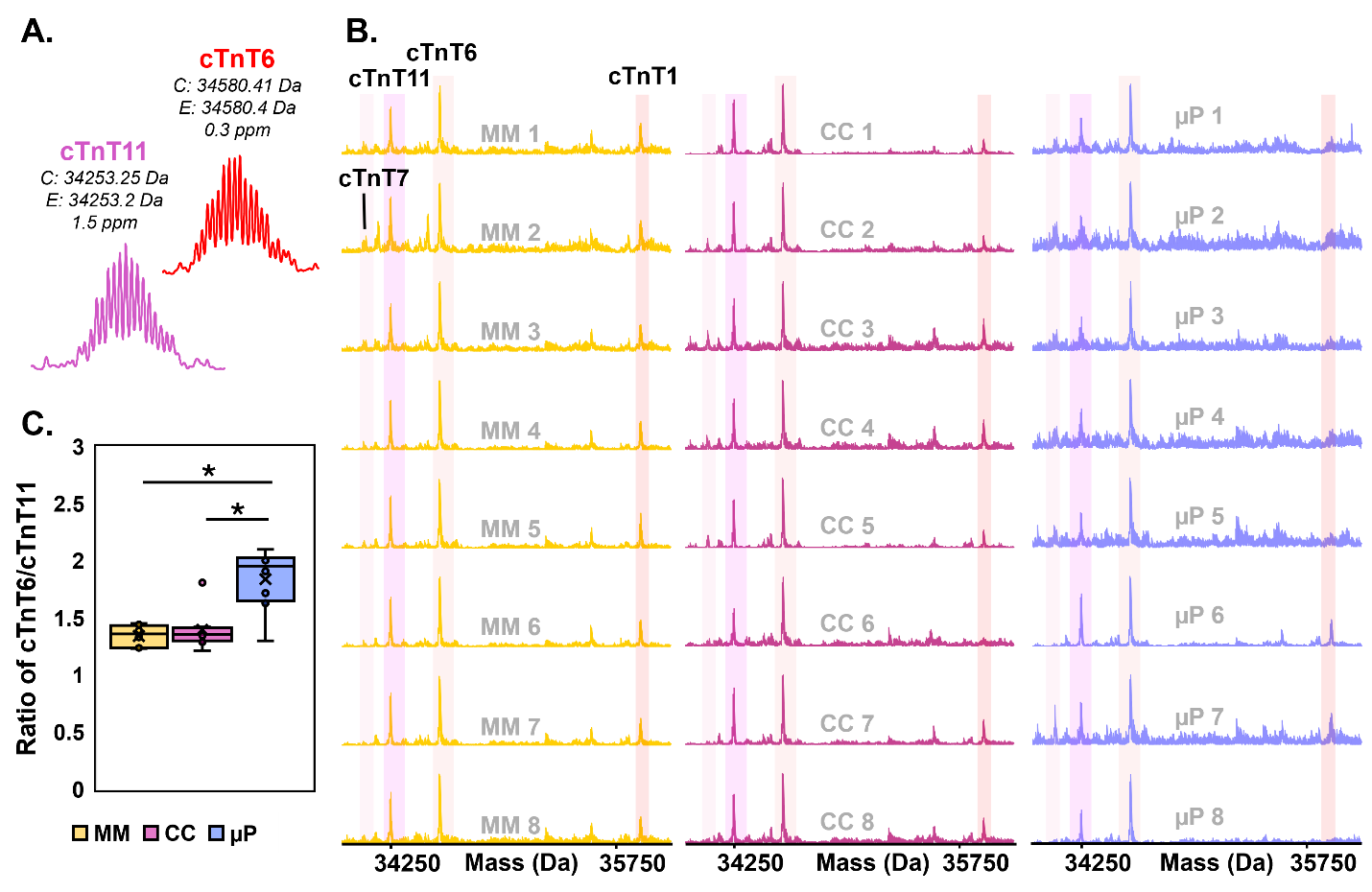

**Figure S8. LC-MS characterization of troponin T isoforms.** A) Representations of the isotopic resolution of troponin T canonical (cTnT6) and non-canonical (cTnT11) isoforms with the calculated and experimental most abundant masses. B) Deconvoluted spectra of cTnT isoforms in each sample across cohorts. Because cTnT isoforms co-elute and have unreliable AUCs, peak intensities were used for the comparison ratio. C) Box plot displaying the ratio of cTnT6 to cTnT11, where “n.s.” indicates a *p* > 0.05, * indicates *p* < 0.05, and ** indicates a *p* < 0.001.

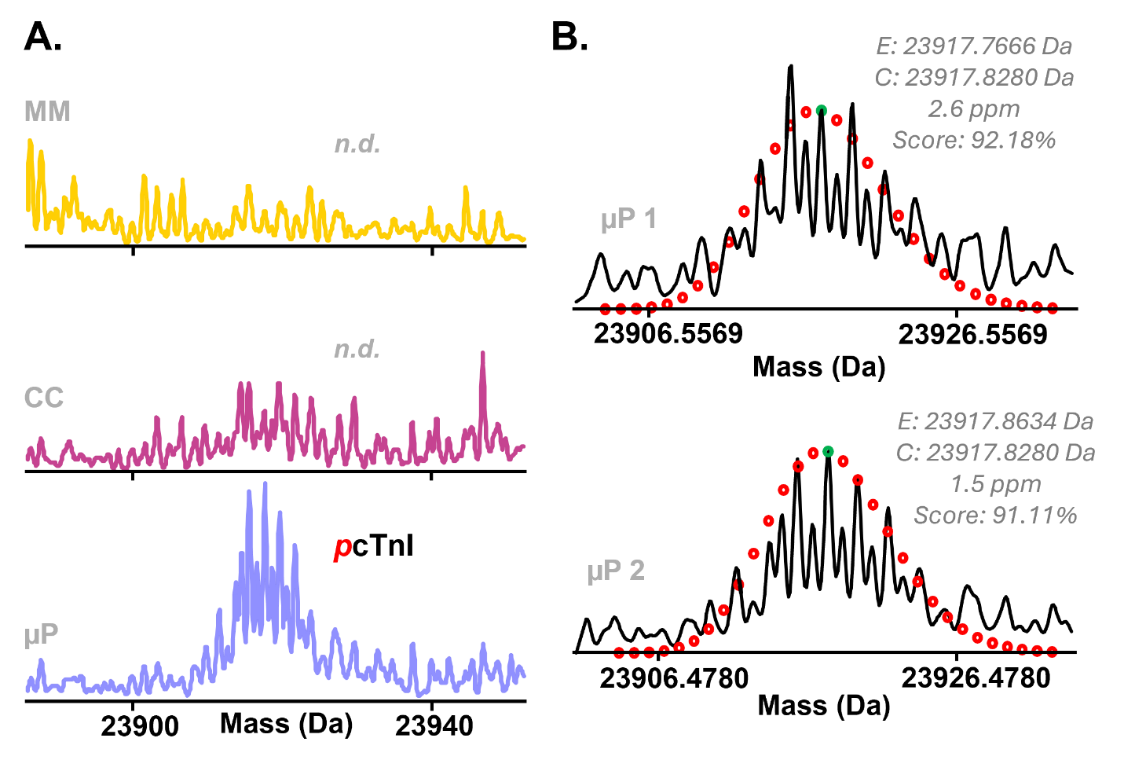

**Figure S9. Identification of low abundance adult cardiac troponin I (cTnI) by top-down LC-MS.** A) Representative deconvoluted mass spectra of MM, CC, and µP displaying the detection of adult cTnI (monophosphorylated) in the µP cohort only. B) Curve-fit regression analysis of cTnI detected in two µP samples (labelled µP 1 and µP 2) enabled by MASH Native showing calculated mass (C), experimental mass (E), error in parts per million (ppm), and the Gaussian distribution expressed as red dots on the deconvoluted spectra.

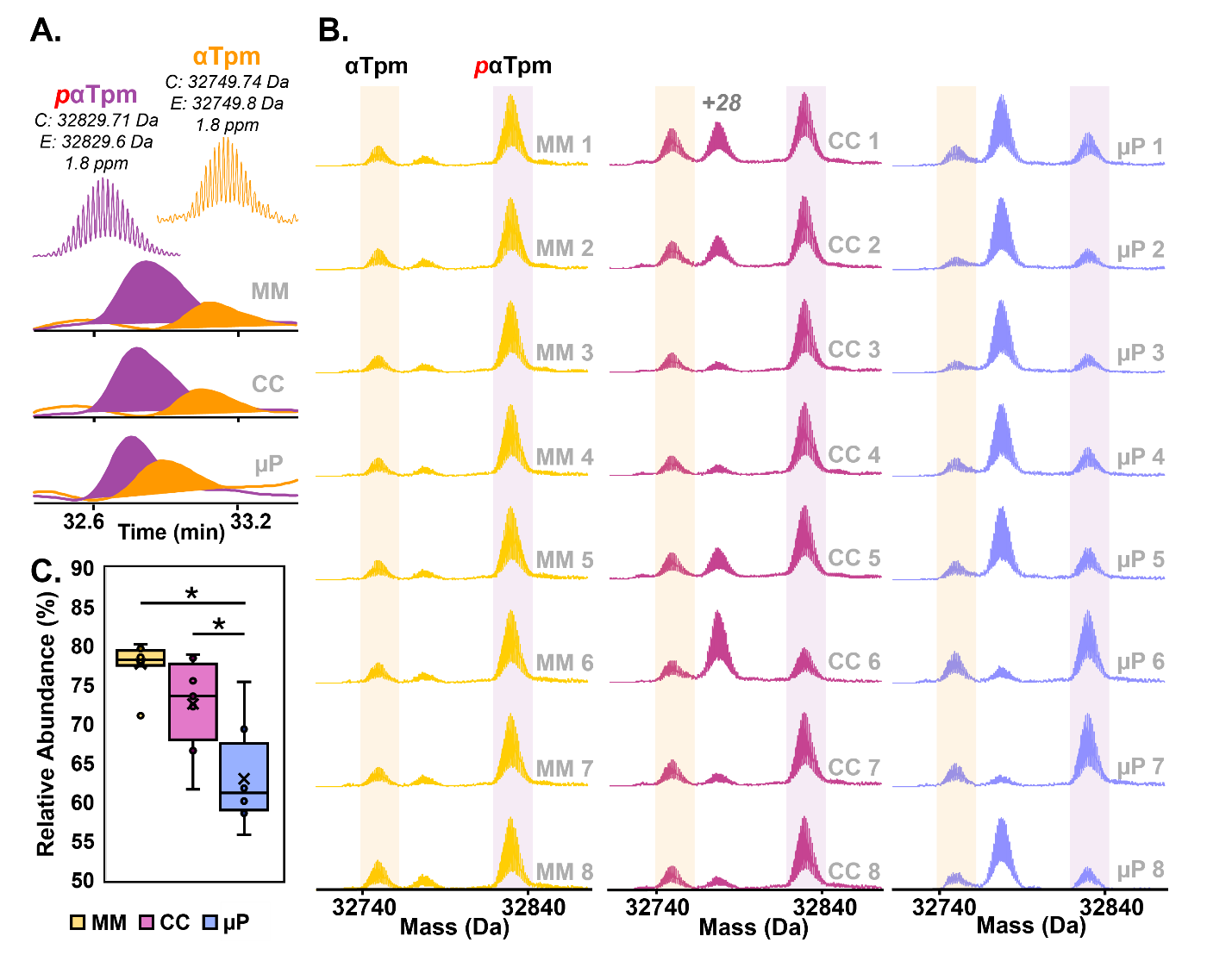

**Figure S10. LC-MS characterization of alpha-Tropomyosin phosphorylation.** A) Representative EICs of MM, CC, and µP cohorts displaying αTpm and its phosphorylated proteoform, where the shaded portion indicates the AUC. Above each, there is a color-matched representation of the isotopic resolution of each isoform with the calculated and experimental most abundant masses. B) Deconvoluted spectra of αTpm and *p*αTpm in each sample across cohorts, which was used for relative abundance quantitation. C) Box plot displaying the relative abundance of *p*αTpm, where “n.s.” indicates a *p* > 0.05, * indicates *p* < 0.05, and ** indicates a *p* < 0.001.

***References***

(1) Lian, X.; Zhang, J.; Azarin, S. M.; Zhu, K.; Hazeltine, L. B.; Bao, X.; Hsiao, C.; Kamp, T. J.; Palecek, S. P. Directed Cardiomyocyte Differentiation from Human Pluripotent Stem Cells by Modulating Wnt/β-Catenin Signaling under Fully Defined Conditions. *Nat Protoc* **2013**, *8* (1), 162–175. <https://doi.org/10.1038/nprot.2012.150>.
(2) Zhang, J.; Tao, R.; Campbell, K. F.; Carvalho, J. L.; Ruiz, E. C.; Kim, G. C.; Schmuck, E. G.; Raval, A. N.; da Rocha, A. M.; Herron, T. J.; Jalife, J.; Thomson, J. A.; Kamp, T. J. Functional Cardiac Fibroblasts Derived from Human Pluripotent Stem Cells via Second Heart Field Progenitors. *Nat Commun* **2019**, *10* (1), 2238. <https://doi.org/10.1038/s41467-019-09831-5>.
(3) Palchesko, R. N.; Zhang, L.; Sun, Y.; Feinberg, A. W. Development of Polydimethylsiloxane Substrates with Tunable Elastic Modulus to Study Cell Mechanobiology in Muscle and Nerve. *PLOS ONE* **2012**, *7* (12), e51499. <https://doi.org/10.1371/journal.pone.0051499>.
(4) Napiwocki, B. N.; Salick, M. R.; Ashton, R. S.; Crone, W. C. Controlling hESC-CM Cell Morphology on Patterned Substrates Over a Range of Stiffness. In *Mechanics of Biological Systems and Materials, Volume 6*; Korach, C. S., Tekalur, S. A., Zavattieri, P., Eds.; Springer International Publishing: Cham, 2017; pp 161–168. <https://doi.org/10.1007/978-3-319-41351-8_23>
(5) Josvai, M.; Polyak, E.; Kalluri, M.; Robertson, S.; Crone, W. C.; Suzuki, M. An Engineered in Vitro Model of the Human Myotendinous Junction. *Acta Biomater* **2024**, *180*, 279–294. <https://doi.org/10.1016/j.actbio.2024.04.007>.
(6) Salick, M. R.; Napiwocki, B. N.; Sha, J.; Knight, G. T.; Chindhy, S. A.; Kamp, T. J.; Ashton, R. S.; Crone, W. C. Micropattern Width Dependent Sarcomere Development in Human ESC-Derived Cardiomyocytes. *Biomaterials* **2014**, *35* (15), 4454–4464. <https://doi.org/10.1016/j.biomaterials.2014.02.001>.
(7) Stempien, A.; Josvai, M.; de Lange, W. J.; Hernandez, J. J.; Notbohm, J.; Kamp, T. J.; Valdivia, H. H.; Eckhardt, L. L.; Maginot, K. R.; Ralphe, J. C.; Crone, W. C. Identifying Features of Cardiac Disease Phenotypes Based on Mechanical Function in a Catecholaminergic Polymorphic Ventricular Tachycardia Model. *Front. Bioeng. Biotechnol.* **2022**, *10*. <https://doi.org/10.3389/fbioe.2022.873531>.
(8) Yu, H.; Xiong, S.; Tay, C. Y.; Leong, W. S.; Tan, L. P. A Novel and Simple Microcontact Printing Technique for Tacky, Soft Substrates and/or Complex Surfaces in Soft Tissue Engineering. *Acta Biomater* **2012**, *8* (3), 1267–1272. <https://doi.org/10.1016/j.actbio.2011.09.006>.
(9) Stempien, A.; Josvai, M.; Notbohm, J.; Zhang, J.; Kamp, T. J.; Crone, W. C. Influence of Remodeled ECM and Co-Culture with iPSC-Derived Cardiac Fibroblasts on the Mechanical Function of Micropatterned iPSC-Derived Cardiomyocytes. *Cardiovasc Eng Technol* **2024**, *15* (3), 264–278. <https://doi.org/10.1007/s13239-024-00711-8>.
(10) Melby, J. A.; Brown, K. A.; Gregorich, Z. R.; Roberts, D. S.; Chapman, E. A.; Ehlers, L. E.; Gao, Z.; Larson, E. J.; Jin, Y.; Lopez, J. R.; Hartung, J.; Zhu, Y.; McIlwain, S. J.; Wang, D.; Guo, W.; Diffee, G. M.; Ge, Y. High Sensitivity Top-down Proteomics Captures Single Muscle Cell Heterogeneity in Large Proteoforms. *Proc Natl Acad Sci U S A* **2023**, *120* (19), e2222081120. <https://doi.org/10.1073/pnas.2222081120>.
(11) Larson, E. J.; Pergande, M. R.; Moss, M. E.; Rossler, K. J.; Wenger, R. K.; Krichel, B.; Josyer, H.; Melby, J. A.; Roberts, D. S.; Pike, K.; Shi, Z.; Chan, H.-J.; Knight, B.; Rogers, H. T.; Brown, K. A.; Ong, I. M.; Jeong, K.; Marty, M. T.; McIlwain, S. J.; Ge, Y. MASH Native: A Unified Solution for Native Top-down Proteomics Data Processing. *Bioinformatics* **2023**, *39* (6), btad359. <https://doi.org/10.1093/bioinformatics/btad359>.
